# Diet-induced obesity alters intestinal monocyte-derived and tissue-resident macrophages in female mice independent of TNF

**DOI:** 10.1101/2022.09.27.509732

**Authors:** Jessica A. Breznik, Jennifer Jury, Elena F. Verdú, Deborah M. Sloboda, Dawn M. E. Bowdish

## Abstract

Macrophages are essential for homeostatic maintenance of the anti-inflammatory and tolerogenic intestinal environment, yet monocyte-derived macrophages can promote local inflammation. Pro-inflammatory macrophage accumulation within the intestines may contribute to the development of systemic chronic inflammation and immunometabolic dysfunction in obesity. Using a model of high fat diet-induced obesity in C57BL/6J female mice, we assessed intestinal permeability by *in vitro* and *in vivo* assays, and quantitated intestinal macrophages in ileum and colon tissues by multicolour flow cytometry after short (6 weeks), intermediate (12 weeks), and prolonged (18 weeks) diet allocation. We characterized monocyte-derived CD4^−^TIM4^−^ and CD4^+^TIM4^−^ macrophages, as well as tissue-resident CD4^+^TIM4^+^ macrophages. Diet-induced obesity had tissue and time-dependent effects on intestinal permeability, as well as monocyte and macrophage numbers, surface phenotype, and intracellular production of the cytokines IL-10 and TNF. We found that obese mice had increased paracellular permeability, in particular within the ileum, but this did not elicit recruitment of monocytes, nor a local pro-inflammatory response by monocyte-derived or tissue-resident macrophages, in either the ileum or colon. Proliferation of monocyte-derived and tissue-resident macrophages was also unchanged. Wildtype and TNF^−/-^ littermate mice had similar intestinal permeability and macrophage population characteristics in response to diet-induced obesity. These data are unique from reported effects of diet-induced obesity on macrophages in metabolic tissues, as well as outcomes of acute inflammation within the intestines, and collectively indicate that TNF does not mediate effects of diet-induced obesity on intestinal monocyte-derived and tissue-resident intestinal macrophages in young female mice.

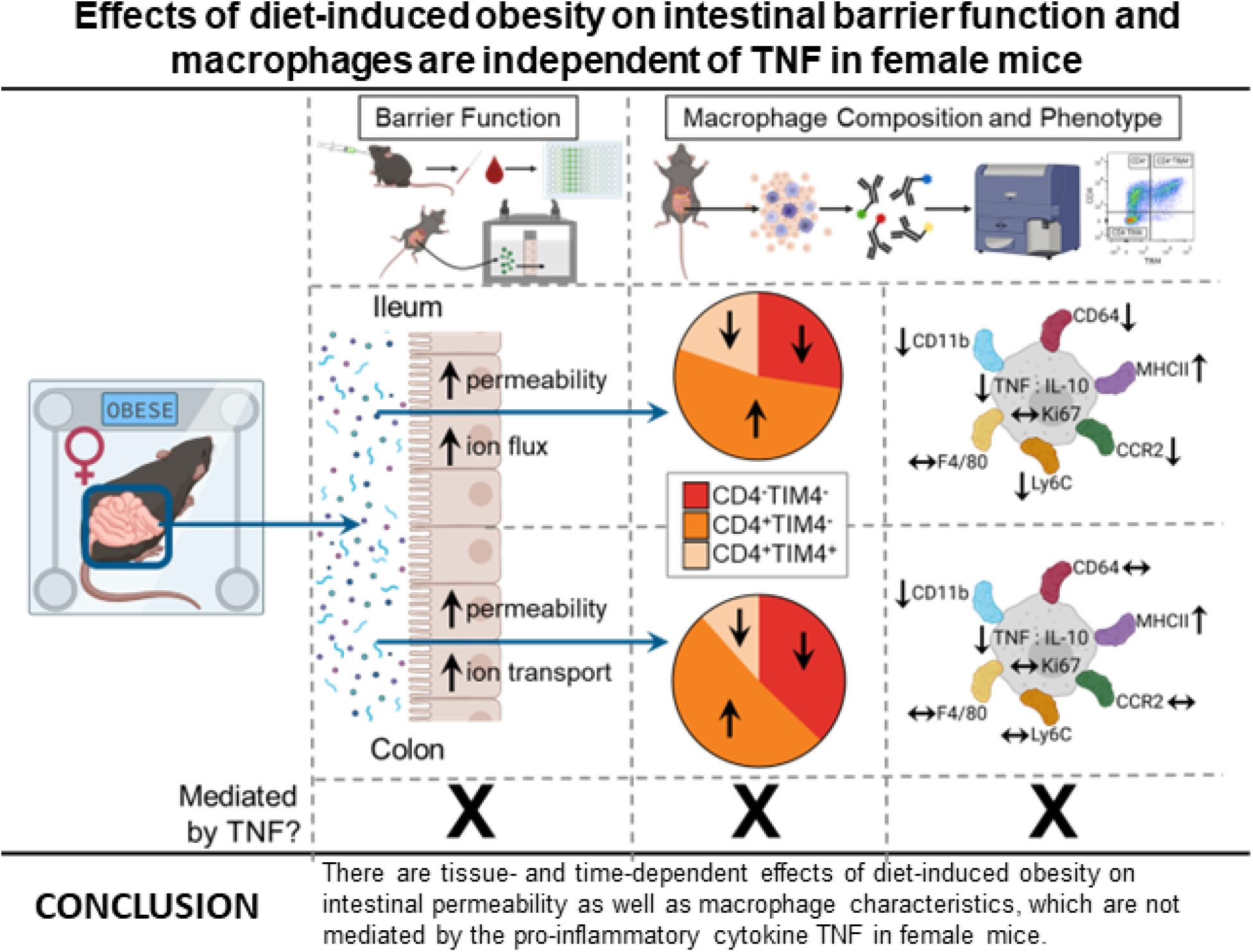

## Introduction

Macrophages are innate immune cells with diverse tissue-specific roles in maintenance of homeostasis (Gordon & Plüddemann, 2019), and in the initiation and resolution of the acute inflammatory response (Medzhitov, 2008). Macrophages also contribute to the pathophysiology of chronic inflammatory diseases such as obesity (McNelis & Olefsky, 2014). Like most tissues, the intestines contain a significant proportion of long-lived tissue-resident macrophages, as well as macrophages derived from blood monocytes, which originate from bone marrow progenitors (De Schepper *et al*., 2018; Shaw *et al*., 2018; Liu *et al*., 2019). Macrophages are present along the entire length and within all structural layers of the intestines (Hume *et al*., 1984; Mikkelsen *et al*., 1988; De Schepper *et al*., 2018). Monocyte-derived macrophages are located proximal to the intestinal epithelium where they maintain epithelial barrier function by mediating interactions between the microbiota, luminal antigens, and other immune cells, to ultimately support immunological tolerance and host defense (De Schepper *et al*., 2018; Shaw *et al*., 2018). Longer-lived tissue-resident macrophages, in contrast, localize in the submucosal and muscularis layers of the intestinal wall and support the functions of blood and lymphatic vessels, enteric neurons, and smooth muscle cells (Gabanyi *et al*., 2016; De Schepper *et al*., 2018; Shaw *et al*., 2018). Monocyte-derived and tissue-resident intestinal macrophages accordingly have distinct responses to inflammation.

Acute intestinal inflammation, as observed in infection, colitis, and after sterile injury, is often accompanied by rapid recruitment of bone marrow-derived Ly6C^high^ monocytes, which develop into immature pro-inflammatory macrophages that contribute to damage of the intestinal epithelial barrier (Zigmond *et al*., 2012; Bain *et al*., 2013; Bain *et al*., 2018). In contrast, tissue-resident macrophages maintain their anti-inflammatory phenotype and functions, preventing tissue injury from inflammation-elicited immune cells (Qualls *et al*., 2006; Weber *et al*., 2011; Bain *et al*., 2013). Changes to monocyte-derived macrophage phenotype and function may also occur within intestinal tissues under conditions of the systemic, low-grade and chronic inflammation that is characteristic of obesity (Hotamisligil, 2017). Obesity increases circulating Ly6C^high^ monocytes (Breznik *et al*., 2018; Breznik *et al*., 2021a), which differentiate into pro-inflammatory macrophages within metabolic tissues like adipose (Weisberg *et al*., 2003; Kanda *et al*., 2006). While it has been reported that there is an accumulation of pro-inflammatory macrophages within the colon in obesity (Kawano *et al*., 2016), which could contribute to observations of obesity-associated increases in gut permeability (Cani *et al*., 2007; Cani *et al*., 2008; Johnson *et al*., 2015; Kawano *et al*., 2016), there is a lack of consensus across published research (Garidou *et al*., 2015; Johnson *et al*., 2015; Hong *et al*., 2017; Luck *et al*., 2019; Rohm *et al*., 2022). This is likely due to differences in the methods and tissue regions of assessment, length of diet allocation, and selection of appropriate surface antigens to identify macrophages as well as to distinguish their monocyte-derived or tissue-resident characteristics. Importantly, it is also unclear which upstream driver(s) in obesity may contribute to changes in intestinal macrophages.

One such driver may be the pro-inflammatory cytokine tumor necrosis factor (TNF). Within the intestines, TNF is constitutively produced at low levels by intestinal epithelial enterocytes, stromal cells, and by macrophages and other immune cells (Leppkes *et al*., 2014; Delgado & Brunner, 2019). TNF regulates monocyte and macrophage survival and function (Parameswaran & Patial, 2010), as well as homeostatic maintenance and repair of the intestinal epithelium (Delgado & Brunner, 2019). However, macrophage overproduction of TNF promotes intestinal inflammation in mouse models of colitis (Varol *et al*., 2009; Laroui *et al*., 2014), and in inflammatory bowel disease in humans (Kamada *et al*., 2008; Lissner *et al*., 2015). Elevated levels of TNF can also mediate disruption of the intestinal barrier (Leppkes *et al*., 2014; Delgado & Brunner, 2019). TNF has in addition been reported to contribute to chronic intestinal inflammation in obesity (Ding *et al*., 2010; Lam *et al*., 2012; Liu *et al*., 2012; Kawano *et al*., 2016). Recruitment of monocytes that differentiate into TNF-producing pro-inflammatory macrophages within intestinal tissues may therefore promote loss of barrier function that contributes to local inflammation as well as systemic immunometabolic dysfunction in obesity (Kawano *et al*., 2016; Hong *et al*., 2017).

In this study, we used a high fat diet model in littermate wildtype and TNF knockout female mice to examine effects of obesity and the role of TNF within both ileum and colon tissues. We assessed intestinal permeability by *in vivo* and *in vitro* methods, and characterized monocyte-derived and tissue-resident intestinal macrophages by flow cytometry after short, intermediate, and prolonged periods of diet allocation, examining their numbers, prevalence, proliferation, surface phenotype, and cytokine profiles. We hypothesized, following from our prior observations of circulating monocytes and adipose tissue macrophages in female mice (Breznik *et al*., 2021a), that there would be an increase in pro-inflammatory TNF-producing monocyte-derived intestinal macrophages with diet-induced obesity, but genetic ablation of TNF would not prevent obesity-induced loss of intestinal barrier function or changes to intestinal macrophage populations.

## Materials and Methods

### Ethical approval

All animal experiments were approved by McMaster University’s Animal Research Ethics Board following the recommendations of the Canadian Council on Animal Care.

### Animals

All experiments in this study used virgin female mice. Mice were originally purchased from The Jackson Laboratory. Wildtype C57BL/6J mice (cat#000664) and tumour necrosis factor knockout (TNF^−/-^) C57BL/6J mice (cat#003008) were bred in-house at the McMaster University Central Animal Facility under specific pathogen free conditions. Littermate TNF^+/+^ and TNF^−/-^ mice were generated from F2 TNF^+/-^ heterozygotes (Robertson *et al*., 2019), with confirmation of genotype by PCR according to standard protocols of The Jackson Laboratory. WT mice were weaned onto a standard chow diet (cat#8640 Teklad 22/5 Rodent Diet, Envigo). Littermate WT and TNF^−/-^ mice were weaned onto Teklad irradiated global 14% protein diet (cat#2914, Envigo). To assess effects of diet-induced obesity, beginning at 5 weeks of age weight-matched wildtype mice were fed *ad libitum* a standard chow diet (17% kcal fat, 29% kcal protein, 54% kcal carbohydrates; Teklad 22/5 Rodent Diet) or a high fat (HF) diet (60% kcal fat, 20% kcal protein, 20% kcal carbohydrates; D12492, Research Diets, Inc.) and provided water *ad libitum*. To assess effects of TNF in diet-induced obesity, beginning at 8 weeks of age littermate TNF^+/+^ and TNF^−/-^ mice were fed *ad libitum* the high fat diet and provided water *ad libitum*. Mice were cohoused 2-5 per cage with constant ambient temperature (22°C) on a 12-hour light-dark cycle under specific pathogen-free conditions. Mice were housed in vent/rack cages with a plastic tube and cotton and paper bedding material for enrichment. Mice were sacrificed by exsanguination and/or cervical dislocation under isoflurane anesthesia.

### *In vivo* intestinal paracellular permeability assay

Mice were placed in a clean cage and fasted for 6 hours (3 am – 9 am) prior to and during the permeability assay. Blood was collected via tail nick using a heparinized capillary tube prior to gavage (baseline or 0 minute sample) with 4 kDa fluorescein isothiocyanate-conjugated dextran (FITC-dextran; cat#46944, Sigma-Aldrich) diluted in PBS (pH 7.4) (80 mg/mL and administered at 500 mg/kg body weight), and again at 30, 60, 90, 120, and/or 240 minutes post-gavage. Acid-citrate dextrose (15% v/w; cat#C3821, Sigma-Aldrich) was added to blood samples after collection to prevent clotting. Plasma was collected post-centrifugation (8000 rpm, 10 minutes) after each time point and stored at 4°C until all samples were collected. Fluorescence was measured in plasma diluted 1:10 in PBS, in duplicate, on a plate reader with excitation at 585 nm and emission at 530 nm (Synergy H4 Hybrid Microplate Reader, BioTek Instruments, Inc.). Whole-intestine permeability at each time point in each mouse was assessed by subtracting the average relative fluorescence units of baseline plasma and triplicate wells of PBS (sample blank) from the average post-gavage relative fluorescence units.

### *In vitro* intestinal paracellular permeability assay

Tissues from the proximal colon and distal ileum were excised, opened along the mesenteric border, stripped of the external muscularis layer, and mounted into Ussing chambers (Physiologic Instruments, Inc.), as previously described (Pinier *et al*., 2012; Silva *et al*., 2012). In brief, samples were equilibrated in oxygenated Krebs buffer containing 10 mM glucose (serosal side) or 10 mM mannitol (luminal side) at 37°C for ∼30 minutes. Short circuit current (Isc) was measured to assess active ion transport. Measurements of potential difference and short circuit current were used to calculate tissue conductance (G; mS/cm^2^) by Ohm’s law. Paracellular permeability was determined by measuring mucosal-to-serosal flux (%flux/cm^2^/hour) of the small inert probe ^51^Cr-EDTA (360 Da; 6 μCi/mL) (Perkin Elmer), by taking an initial radiolabelled sample of mucosal buffer and then samples from the serosal compartment every 30 minutes for 2 hours, with addition of fresh buffer to maintain a constant volume. Samples were assessed in a liquid scintillation counter (LS6500 Multi Purpose Scintillation Counter, Beckman Coulter, Inc.), with radioactive counts from each 30 minute time point averaged and compared to the initial radiolabelled sample.

### Processing of intestinal tissues for flow cytometry

Intestinal ileum and colon lengths were measured, and the tissues were processed for flow cytometry based on previous protocols (Sun *et al*., 2007; Shaw *et al*., 2018). The distal 25% of the small intestine adjacent to the caecum was considered ileal tissue. After dissection, intestinal tissue was immediately placed in ice-cold PBS. Peyer’s patches (in the small intestine) and mesenteric fat were removed, and the tissue was cut longitudinally and washed with PBS to remove contents. The tissue was cut into ∼2 cm pieces and incubated in pre-warmed ‘Stir Media’ (RPMI 1640 supplemented with 3% FBS, 100 U/mL penicillin, 100 μg/mL streptomycin, 20 mM HEPES, 5 mM EDTA pH 8.0, and 1 mM DTT (cat#D0632, Sigma-Aldrich)) at 37°C for 20 minutes with constant stirring (∼550 rpm) to remove the mucus layer. Tissues were washed 3x by shaking for 30 seconds in serum-free pre-warmed RPMI 1640 supplemented with 100 U/mL penicillin, 100 μg/mL streptomycin, 2 mM EDTA, and 20 mM HEPES, and rinsed with pre-warmed PBS. The tissue was finely minced with scissors and incubated by stirring in serum-free pre-warmed ‘Complete Media’ (RPMI 1640 supplemented with 100 U/mL penicillin, 100 μg/mL streptomycin, 20 mM HEPES, 1% (v/v) non-essential amino acids, 1% (v/v) sodium pyruvate, 1% (v/v) L-glutamine, 0.01% (v/v) β-mercaptoethanol), with 0.5 mg/mL DNAse I (cat#10104159001, Sigma-Aldrich) and 0.1 mg/mL Liberase TL (cat#05401020001, Sigma- Aldrich) added immediately prior to incubation for 30 minutes at 37°C (stirring at ∼550 rpm), to dissociate intestinal epithelial cells and leukocytes. Digested tissue was homogenized, washed with ice-cold Stir Media, passed through a 70 μm filter, centrifuged at 1500 rpm for 10 minutes, resuspended in ice-cold Stir Media, passed through a 40 μm filter, and centrifuged again at 1500 rpm for 10 minutes. The pellet was resuspended in ice-cold Complete Media supplemented with 3% fetal calf serum until staining.

### Flow cytometry analysis

Live cells were counted manually under a light microscope with a hemocytometer and trypan blue (cat#MT25900CI, Corning). Single cell suspensions of 5 × 10^5^ – 2 × 10^6^ cells were stained in a 96 v-well plate, with all centrifugation steps at 2000 rpm for 2 minutes. Samples were washed in PBS and were incubated at 4°C with CD16/32 Fc block (cat#101302, BioLegend) in PBS for 10 minutes before staining with antibodies (Supplementary Table 1) in PBS for 20 minutes at 4°C. Samples were washed twice in PBS, fixed for 10 minutes in 1x eBioscience Fix-Lyse buffer (cat#00-5333-52, ThermoFisher Scientific), washed in PBS, and resuspended in PBS for flow cytometer analysis. For intracellular staining without stimulation, samples were stained as described above with a modified surface stain (Supplementary Table 2), and then permeabilized in 1x eBioscience Permeabilization Buffer (cat#88-8824-00, ThermoFisher Scientific) for 30 minutes at room temperature, stained with intracellular antibody mix (Supplementary Table 3) in 1x Permeabilization Buffer for 30 minutes, washed twice in PBS, and resuspended in PBS for flow cytometer analysis.

Samples were run on a BD Biosciences Fortessa flow cytometer (BD Biosciences). Data were analyzed using the FlowJo v9 software (Tree Star). Stained cells were assessed with unstained, isotype, and/or fluorescence-minus-one controls. Expression of surface markers was quantified by measuring geometric mean fluorescence intensity of each fluorescence marker and subtracting background geometric mean fluorescence intensity of isotype or fluorescence-minus-one controls. Geometric mean expression of each surface marker was combined across multiple independent experiments where noted by normalizing the data. Data was normalized by dividing each geometric mean fluorescence data point by the mean of the comparator group (i.e., chow-fed mice in diet experiments and baseline mice in littermate experiments) in each independent experiment. Absolute cell counts (i.e., total cell numbers) were determined with CountBright Absolute Counting Beads (cat#C36950, Life Technologies). Flow cytometry data were analyzed as illustrated in Supplementary Figure 1 for surface staining of intestinal macrophages, and as shown in Supplementary Figure 2 for intracellular staining of intestinal macrophages.

### Statistical analysis

Data were analyzed and plotted with GraphPad Prism version 9 (GraphPad Software). Two-group comparisons of leukocyte populations between diet groups or genotypes were analyzed by unpaired two-tailed Student’s t test (parametric) with Welch’s correction (for unequal variances) or Mann-Whitney U test (non-parametric) according to normality. Comparisons of body weight and intestinal lengths across multiple time points were performed by two-way ANOVA with correction for multiple comparisons by Tukey’s test.

## Results

### Diet-induced obesity decreases intestinal length and increases permeability in female mice

Female mice were fed a standard chow diet or high fat diet and assessed after short (6 weeks), intermediate (12 weeks), and prolonged (18 weeks) diet consumption, based on turnover rates of intestinal macrophages (De Schepper *et al*., 2018; Shaw *et al*., 2018; Liu *et al*., 2019). HF-fed mice gained more weight at each time point compared to chow-fed mice (Figure 1A). Decreased intestinal length has often been considered to be an indication of inflammation, and is frequently observed in mouse models of colitis (Yan *et al*., 2009) and diet-induced obesity (de Wit *et al*., 2008; Ding *et al*., 2010; Navarrete *et al*., 2015; Kawano *et al*., 2016). We observed that after short term diet intake (at 6 weeks) small intestine length was decreased in mice fed HF diet (mean ± SD: 31.6 ± 1.6 cm) compared to chow-fed mice (35.5 ± 2.8 cm), and this effect persisted with intermediate and prolonged diet allocation (i.e. at 12 and 18 weeks) (Figure 1B). Colon length was also significantly decreased in HF-fed mice after 6 weeks (mean ± SD: 6.2 ± 0.4 cm) compared to mice fed chow diet (7.6 ± 1.2 cm) and remained shorter in mice fed HF diet after intermediate and prolonged diet allocation (Figure 1C).

**Figure 1.**
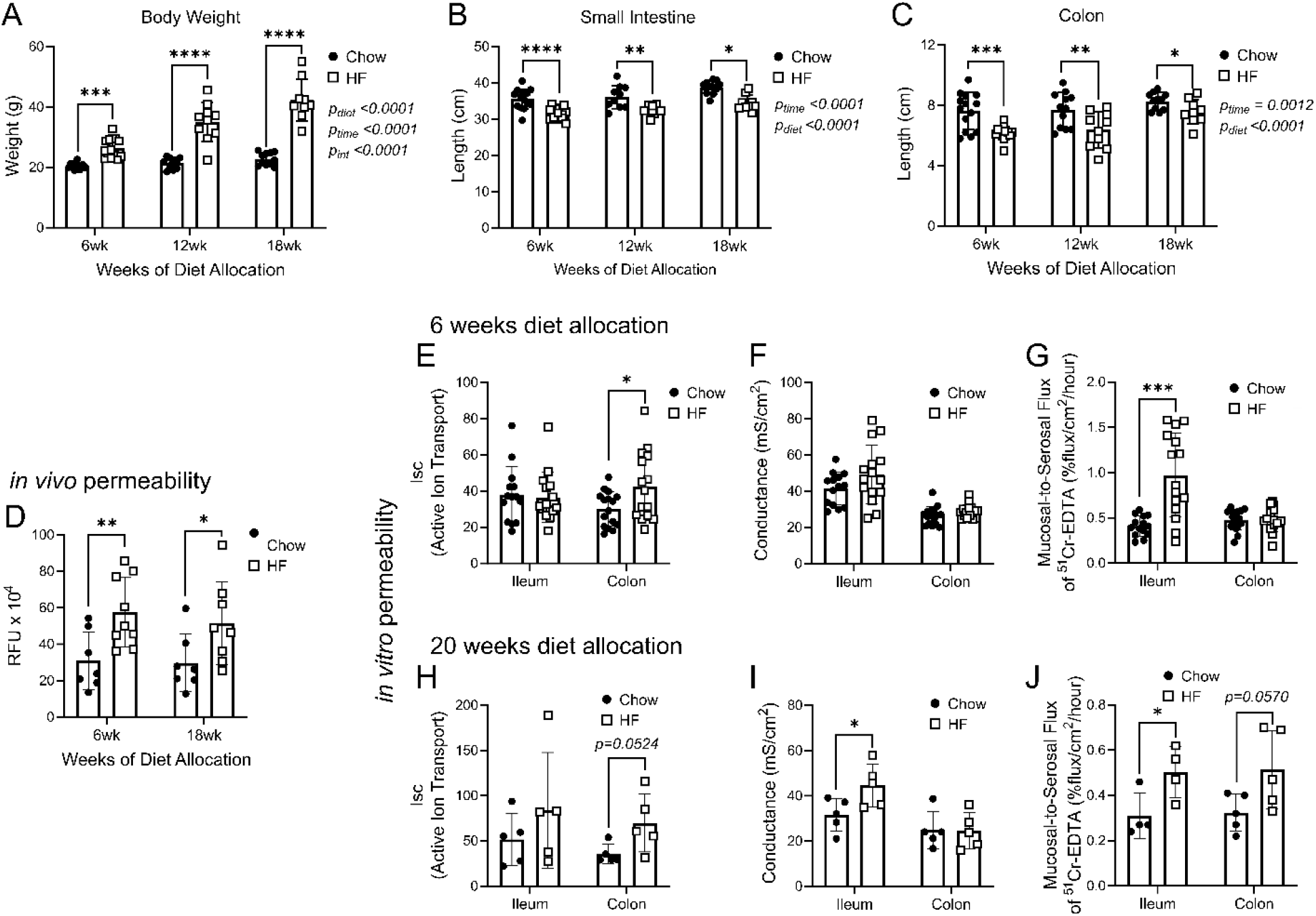
Diet-induced obesity decreases intestinal lengths and increases paracellular permeability in female mice. Body weight, intestinal lengths and permeability were assessed in mice fed a standard chow (Chow) or high fat (HF) diet. (A) body weight and lengths of the (B) small intestine and (C) colon after 6-, 12-, and 18-weeks diet allocation. (D) after 6 and 18 weeks of diet allocation, *in vivo* intestinal permeability to FITC-dextran was measured 4 hours after gavage. *In vitro* ileum and colon tissue active ion transport (E, H), conductance (F, I), and paracellular permeability (G, J) after 6- and 20-weeks diet allocation. Each data point indicates an individual mouse. Data are presented with box height at the mean with error bars indicating ± standard deviation. Data in A-C are combined from two or three independent experiments of n=4-5 mice per group (n=9-14 per time point). Data in D are representative of two independent experiments of n=7-10 mice per diet group and are presented as relative fluorescence units (RFU). Data in E-G (6 weeks) are combined from two independent experiments of n=8 mice per group. Data in H-J (20 weeks) are from one independent experiment of n=4-5 mice per group. Statistical significance was assessed by two-way ANOVA with Tukey’s post-hoc test for between-diet comparisons for A-C and by two-tailed Student’s t test or Welch’s t test for unequal variances for D-J. *\*p<0*.*05, **p<0*.*01, ***p<0*.*001, ****p<0*.*0001*.

To assess if diet-induced obesity affected intestinal permeability in female mice, we initially performed an *in vivo* assay by measuring plasma FITC fluorescence after oral gavage of FITC-dextran. Whole-intestine paracellular permeability increased in HF-fed mice compared to chow-fed mice with short-term (6 weeks) diet allocation, and this increase persisted after prolonged (18 weeks) HF diet intake (Figure 1D). We subsequently used an *in vitro* short-term organ culture method to examine local tissue permeability in the distal ileum and proximal colon after short (6 weeks; Figure 1E-G) and prolonged (20 weeks; Figure 1H-J) HF diet intake. We measured short-circuit current (Isc; i.e., total ion transport as an indication of fluid and electrolyte homeostasis), conductance (i.e., ion flux in relation to tight junction function), and mucosal-to-serosal flux of ^51^Cr-EDTA (i.e., paracellular permeability as an indication of tight junction function) (Clarke, 2009; Thomson *et al*., 2019). Short-circuit current was similar between diet groups after short and prolonged diet allocation in the ileum (Figure 1E, H). Conductance was similar in chow-fed and HF-fed mouse ileum tissues after short-term diet intake (Figure 1F) but increased in HF-fed mouse ileum tissues with longer diet allocation (Figure 1I). Paracellular uptake of ^51^Cr-EDTA significantly increased in ileum tissues of HF-fed mice compared to chow-fed mice at 6 weeks diet intake (Figure 1G) and remained elevated at 20 weeks (Figure 1J). In colon tissues of HF-fed mice compared to chow-fed mice, short-circuit current increased after 6 weeks diet intake (Figure 1E), and a tendency for higher short circuit current was also observed after 20 weeks (Figure 1H), suggestive of a sustained increase in active ion transport and water movement. Colon tissue conductance was similar between diet groups (Figure 1F, I). Paracellular permeability of colon tissues was similar between diet groups after short-term diet allocation (Figure 1G), but tended to be higher after prolonged HF diet intake (Figure 1J). Therefore, both colon and ileum intestinal barrier changes may contribute to the increased total intestinal permeability observed by *in vivo* assay after short and prolonged HF diet. These data collectively indicate that intestinal lengths, active ion transport, and paracellular permeability, are altered within 6 weeks of HF diet consumption. These changes persist or become exacerbated with increasing length of HF diet intake (i.e., after 18-20 weeks).

### Diet-induced obesity alters monocyte and macrophage dynamics in the ileum and colon of female mice

We used flow cytometry to assess monocyte and macrophage populations. We initially quantified Ly6C^high^ monocytes, transitional Ly6C^+^MHCII^+^ cells, and MHCII^+^ macrophages in the ileum (Figure 2A-C) and colon (Figure 2H-J). The numbers of these cells within the ileum were consistently lower after 6-, 12-, and 18-weeks HF diet intake compared to chow-fed mice (Figure 2A-C). Similarly, Ly6C^high^ monocyte and total MHCII^+^ macrophage numbers were lower in colon tissue of HF-fed mice at 6 weeks (Figure 2H and 2J), and Ly6C^high^ monocytes and transitional Ly6C^+^MHCII^+^ cells were lower at 12 weeks (Figure 2H-I), compared to chow-fed mice. In contrast, at a prolonged 18 weeks of diet allocation, all immune cell populations were similar between diet groups in the colon. Since the lengths of the small intestine and colon were decreased in HF-fed mice (Figure 1B-C), we adjusted cell numbers by tissue length and found similar changes (Supplementary Figure 3A-C and 3G-I). These data were also evaluated in terms of relative prevalence (as a proportion of total CD45^+^ leukocytes; Supplementary Figure 3D-F and 3J-L). In the ileum, the relative prevalence of Ly6C^high^ monocytes was significantly lower at all time points in HF-fed dams, whereas the prevalence of MHCII^+^ macrophages was consistently increased, but macrophage prevalence did not increase in the colon. Therefore, while we hypothesized there would be an increase in intestinal monocytes and macrophages with diet-induced obesity, the total numbers of Ly6C^high^ monocytes, transitional Ly6C^+^MHCII^+^ cells, and total MHCII^+^ macrophages, did not increase in the ileum or colon in female mice with diet-induced obesity after short, intermediate, or prolonged HF diet intake.

**Figure 2.**
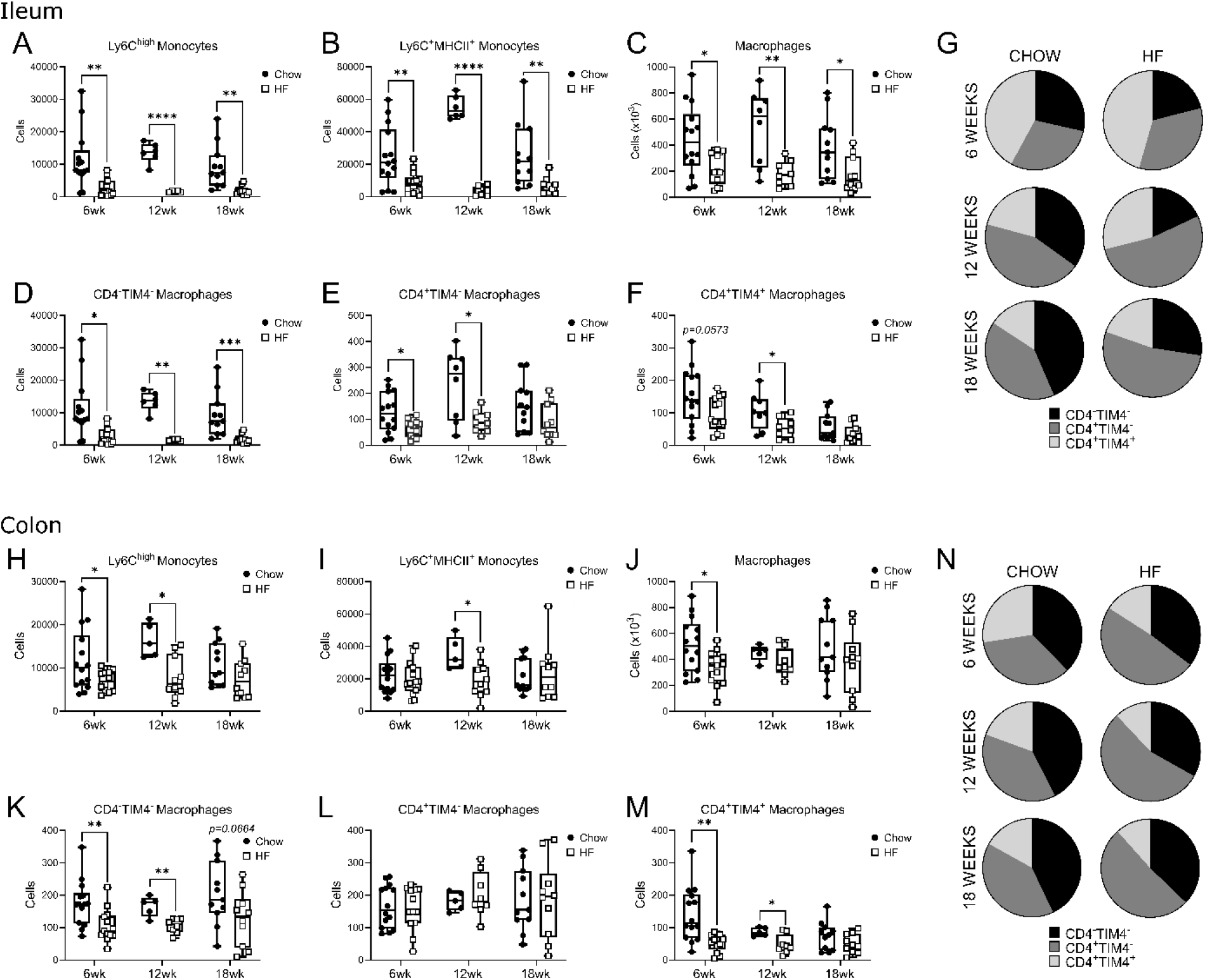
Diet-induced obesity decreases monocytes and macrophages in intestinal tissues in female mice. Intestinal monocyte and macrophage populations were assessed by flow cytometry in the ileum and colon after 6, 12, or 18 weeks of standard chow (Chow) or high fat (HF) diet intake. Ileal absolute cell counts (numbers) of: (A) Ly6C^high^ monocytes, (B) Ly6C^+^MHCII^+^ cells, (C) total MHCII^+^ macrophages, (D) CD4^−^TIM4^−^ macrophages, (E) CD4^+^TIM4^−^ macrophages, (F) CD4^+^TIM4^+^ macrophages. (G) summary of average ileum macrophage prevalence (as a proportion of total macrophages; see Supplementary Figure 4). Colon absolute cell counts (numbers) of: (H) Ly6C^high^ monocytes, (I) Ly6C^+^MHCII^+^ cells, (J) total MHCII^+^ macrophages, (K) CD4^−^TIM4^−^ macrophages, (L) CD4^+^TIM4^−^ macrophages, (M) CD4^+^TIM4^+^ macrophages. (N) summary of average colon macrophage prevalence (as a proportion of total macrophages; see Supplementary Figure 4). Each data point indicates an individual mouse. Data in A-F and H-M are presented as box and whisker plots, minimum to maximum, with the center line at the median. Data are combined from two or three independent experiments of n=4-5 mice per group. Statistical significance was assessed by two-tailed parametric Student’s t test or Welch’s t test for unequal variances or by non-parametric Mann-Whitney U test at each time point. *\*p<0*.*05, **p<0*.*01, ***p<0*.*001, ****p<0*.*0001*.

We next examined the quantity (as total cell numbers and cell numbers per tissue length) and prevalence (as a proportion of total macrophages) of monocyte-derived CD4^−^TIM4^−^ and CD4^+^TIM4^−^ macrophages, as well as tissue-resident CD4^+^TIM4^+^ macrophages, in HF-fed and chow-fed mice after 6-, 12-, and 18-weeks diet intake (Figure 2D-G and K-N; Supplementary Figure 4). Consistent with the observed decreases in ileum Ly6C^high^ monocyte and transitional and Ly6C^+^MHCII^+^ cell populations, monocyte-derived CD4^−^TIM4^−^ macrophage cell numbers (Figure 2D), cell numbers adjusted to tissue length (Supplementary Figure 4A), were lower in ileal tissue of HF-fed mice compared to chow-fed mice, whether after short, intermediate, or prolonged HF diet allocation. Total cell quantities (and cell numbers adjusted by tissue length) of ileal CD4^+^TIM4^−^ macrophage populations were also lower at short and intermediate (i.e. 6 and 12 weeks), but not prolonged (18 weeks), periods of HF diet intake (Figure 2E; Supplementary Figure 4B). Quantities of tissue resident CD4^+^TIM4^+^ macrophages (but not cell numbers adjusted by tissue length) were also lower at 12 but not 18 weeks of HF diet intake (Figure 2F; Supplementary Figure 4C). These changes resulted in a proportional decrease in ileum CD4^−^ TIM4^−^ macrophages at all assessed time points, and an increase in the prevalence of CD4^+^TIM4^−^ macrophages at 12 and 18 weeks (Figure 2G and Supplementary Figure 4D-F). Intra-macrophage population dynamics were also altered in the colon. CD4^+^TIM4^−^ macrophage numbers, and numbers adjusted by tissue length, were similar between diet groups at all time points (Figure 2L; Supplementary Figure 4H). However, CD4^+^TIM4^+^ cell numbers and numbers adjusted by colon tissue length were lower in HF-fed compared to chow-fed mice at 6 and 12 weeks, but not 18 weeks (Figure 2M; Supplementary Figure 4I). These changes resulted in a significantly reduced prevalence of CD4^+^TIM4^+^ macrophages and an increase in CD4^+^TIM4^−^ macrophages (as a proportion of total macrophages) after 6, 12, and 18 weeks of HF diet intake (Figure 2N; Supplementary Figure 4J-L). Therefore, these data show that there are temporal and tissue-specific changes to intestinal monocyte and macrophage dynamics in response to diet-induced obesity.

**Figure 4.**
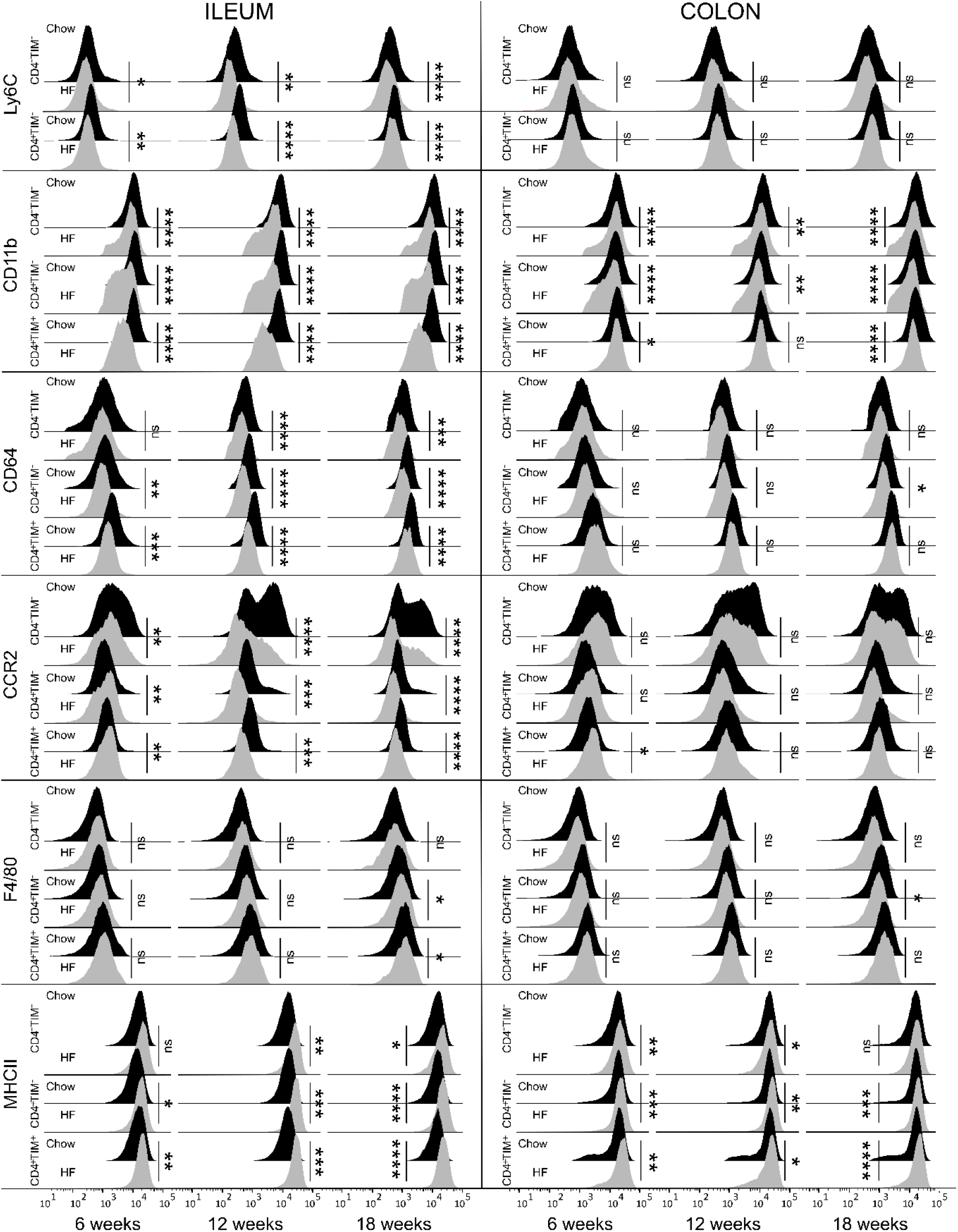
Intestinal macrophage surface phenotype is altered between chow and HF-fed mice. Flow cytometry analysis of CD4^−^TIM4^−^, CD4^+^TIM4^−^, and CD4^+^TIM4^+^ macrophage populations in the ileums and colons of chow and HF-fed female mice after 6-, 12-, or 18-weeks diet allocation. Macrophage phenotype was assessed by examining surface expression of Ly6C, CD11b, CD64, CCR2, F4/80, and MHCII. Data are visualized by concatenating uncompensated events in FlowJo for each mouse and indicated macrophage population at each time point grouped by diet type, and then geometric mean fluorescence intensity expression data of each concatenated group was overlaid onto the same histogram plot. Data are from 2-3 independent experiments of n=4-5 mice per group at each time point. Statistical significance was assessed by two-tailed parametric Student’s t test or Welch’s t test for unequal variances or by non-parametric Mann-Whitney U test at each time point. *\*p<0*.*05, **p<0*.*01, ***p<0*.*001, ****p<0*.*0001*.

### Diet-induced obesity does not alter intestinal macrophage proliferation in female mice, but increases intracellular IL-10 and alters surface phenotype

In addition to reduced monocyte recruitment, a loss of proliferative capacity could contribute to the reduced quantities of macrophages that we observed in the ileum and colon after HF diet intake. We assessed macrophage proliferation in the ileum and colon by flow cytometry via intracellular staining of the nuclear antigen Ki67 (Figure 3A-B,E). Consistent with prior observations (Bain *et al*., 2014), we observed Ki67 expression suggestive of macrophage proliferation. However, there were no statistically significant differences in Ki67 expression between HF and chow diet groups for CD4^−^TIM4^−^, CD4^+^TIM4^−^, or CD4^+^TIM4^+^ macrophages in the ileum or colon at any assessed time point. Thus, HF diet-induced obesity did not appear to alter the self-renewal rate of tissue-resident CD4^+^TIM4^+^ macrophages. Nor was the proliferation of monocyte-derived CD4^−^TIM4^−^ macrophages, or CD4^+^TIM4^−^ macrophages, changed in HF-fed mice compared to chow-fed mice, after short, intermediate, or prolonged HF diet intake.

**Figure 3.**
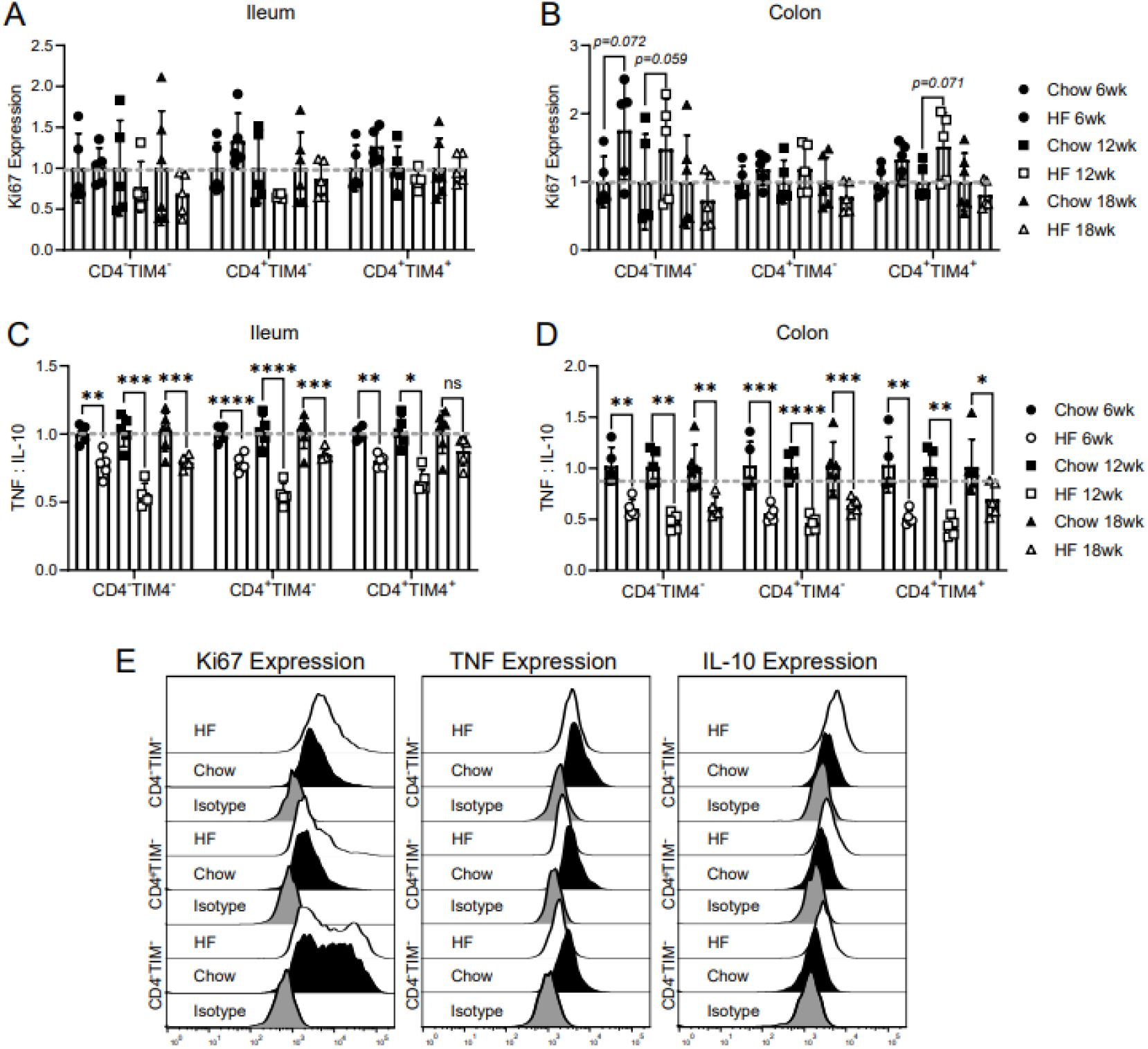
Colon and ileum macrophage proliferation and intracellular production of IL-10 and TNF is altered in obese female mice. Ileum and colon CD4^−^TIM4^−^, CD4^+^TIM4^−^, and CD4^+^TIM4^+^ macrophage proliferation (by Ki67) and intracellular cytokine expression (of TNF and IL-10) was assessed by flow cytometry in standard chow (Chow) or high fat (HF) diet fed female mice at 6, 12, or 18 weeks of diet intake. Proliferation of macrophages after 6, 12, and 18 weeks in the (A) ileum and (B) colon. Ratio of IL-10 to TNF expression of macrophages after 6, 12, and 18 weeks in the (C) ileum and (D) colon. (E) representative expression of Ki67, TNF, and IL-10 in macrophages of chow-fed and HF-fed mice compared to isotype controls. Macrophage proliferation was compared across lengths of diet allocation (i.e., independent experiments) by normalizing the geometric mean data from each mouse to the mean of the chow mouse group for each macrophage population in each independent experiment. Individual cytokine expression measurements of TNF and IL-10 are shown in Supplementary Figure 5. Each data point indicates an individual mouse. Data are from one independent experiment at each time point of n=5-6 mice per group. Data are presented with box height at the mean with error bars at ± standard deviation. Statistical significance was assessed by two-tailed parametric Student’s t test or Welch’s t test for unequal variances or by non-parametric Mann-Whitney U test at each time point. *\*p<0*.*05, **p<0*.*01, ***p<0*.*001, ****p<0*.*0001*.

Constitutive expression of IL-10 by intestinal macrophages under homeostatic conditions supports their roles in maintenance of the intestinal barrier and a tolerogenic intestinal environment (Shouval *et al*., 2014; Morhardt *et al*., 2019). Intestinal macrophages are also a primary source of TNF. Low levels of TNF are necessary for maintenance of the intestinal epithelium, but higher levels are associated with loss of gut homeostasis (Leppkes *et al*., 2014; Delgado & Brunner, 2019). We examined intracellular IL-10 and TNF in intestinal macrophages of chow and HF-fed mice in the absence of exogenous stimulation (Figure 3C-E; Supplementary Figure 5). Consistent with previous studies (Rivollier *et al*., 2012; Bain *et al*., 2013), all intestinal macrophages produced IL-10 as well as TNF, although we found important macrophage population and diet-associated differences. In chow-fed mice, CD4^−^TIM4^−^ macrophages produced more TNF per cell than CD4^+^TIM4^−^ or CD4^+^TIM4^+^ macrophages. There were also temporal changes to IL-10 and TNF expression within all macrophage populations from short, intermediate, and prolonged HF diet intake, but intracellular TNF expression in HF-fed mice was not elevated in CD4^−^TIM4^−^, CD4^+^TIM4^−^ or CD4^+^TIM4^+^ macrophages in either the ileum or the colon. Furthermore, the ratio of TNF to IL-10 intracellular expression was lower in intestinal macrophages from HF-fed mice compared to chow-fed mice.

While our observations of altered intracellular macrophage TNF and IL-10 expression may be suggestive of a more anti-inflammatory role of ileum and colon macrophages after HF diet intake, changes to surface antigen expression can also provide a preliminary assessment of cellular function. We examined ileum and colon macrophage surface expression of Ly6C, CD11b, CD64, CCR2, F4/80, and MHCII (Figure 4). These surface markers are associated with macrophage migration (CCR2, CD11b), activation (CD11b, F4/80, CD64), phagocytosis and antigen presentation (MHCII), as well as maturity (Ly6C, CD64, F4/80, MHCII) (Reith *et al*., 2005; Freeman *et al*., 2010; Han *et al*., 2010; Shi & Pamer, 2011; Tamoutounour *et al*., 2012; Bain *et al*., 2014; Gordon & Plüddemann, 2019). As monocytes differentiate into intestinal tissue macrophages, they progressively lose expression of Ly6C and CCR2 while increasing their expression of CD64, F4/80, and MHCII (Guilliams *et al*., 2020). This assessment revealed significant diet-associated differences in macrophage surface phenotype, particularly within the ileum. Ileal CD4^−^TIM4^−^, CD4^+^TIM4^−^, and CD4^+^TIM4^+^ macrophages had reduced surface expression of Ly6C, CD11b, CD64, and CCR2 after short term (6 weeks) and prolonged (18 weeks) HF diet intake. Compared to chow-fed mice, MHCII expression was higher on all ileum macrophage populations of HF-fed mice within 12 weeks, and remained higher at 18 weeks. As in the ileum, all colon macrophage populations of HF-fed mice compared to chow-fed mice had lower expression of CD11b within 6 weeks of diet intake, which was also apparent after prolonged diet intake at 18 weeks, whereas CD4^+^TIM4^−^ and CD4^+^TIM4^+^ macrophages in HF-fed mice had constitutively higher MHCII expression. Colon CD4^+^TIM4^−^ macrophages also had decreased CD64 expression at 18 weeks. Therefore, short, intermediate and prolonged HF diet intake altered the surface phenotypes of ileum and colon monocyte-derived CD4^−^TIM^−^ and CD4^+^TIM4^−^ macrophages, as well as tissue-resident CD4^+^TIM4^+^ macrophages. Overall, these data demonstrate that diet-induced obesity results in macrophage subset, tissue, and time-dependent effects on intestinal macrophage surface phenotype and intracellular cytokine production.

### TNF^−/-^ female mice are not protected from obesity-induced changes in intestinal length, permeability, or macrophage population dynamics

To assess the involvement of TNF-mediated inflammation on intestinal physiology and macrophages in diet-induced obesity, littermate TNF^−/-^ and wildtype female mice were examined prior to HF diet allocation (i.e. baseline), or after 15 and 30 weeks of HF diet intake. We observed a main effect of time of HF diet allocation on body weight, but body weight was similar between genotypes (Figure 5A). There were main effects of genotype and time on small intestine lengths (Figure 5B). The small intestines of HF-fed WT mice (mean ± SD, 30 weeks: 32.1 ± 1.5 cm) were shorter compared to baseline chow-fed WT mice (baseline: 36.3 ± 0.3 cm), as were the small intestine lengths of HF-fed TNF^−/-^ mice (30 weeks: 33.1 ± 1.4 cm) compared to baseline chow-fed TNF^−/-^ mice (baseline: 37.9 ± 2.2 cm), though post-hoc tests were not significant between genotypes. While there was no significant effect of genotype on colon length, it decreased with increasing time of HF diet allocation (Figure 5C). *In vivo* assessments of paracellular permeability showed no significant differences by genotype after 12 weeks (Figure 5D) or 28 weeks (Figure 5E) of HF diet intake, suggesting that TNF^−/-^ mice had similar whole-intestine permeability as WT mice. Together, these data show that in female mice, TNF is not a primary driver of phenotypic changes to body weight, intestinal length, or paracellular permeability, in diet-induced obesity.

**Figure 5.**
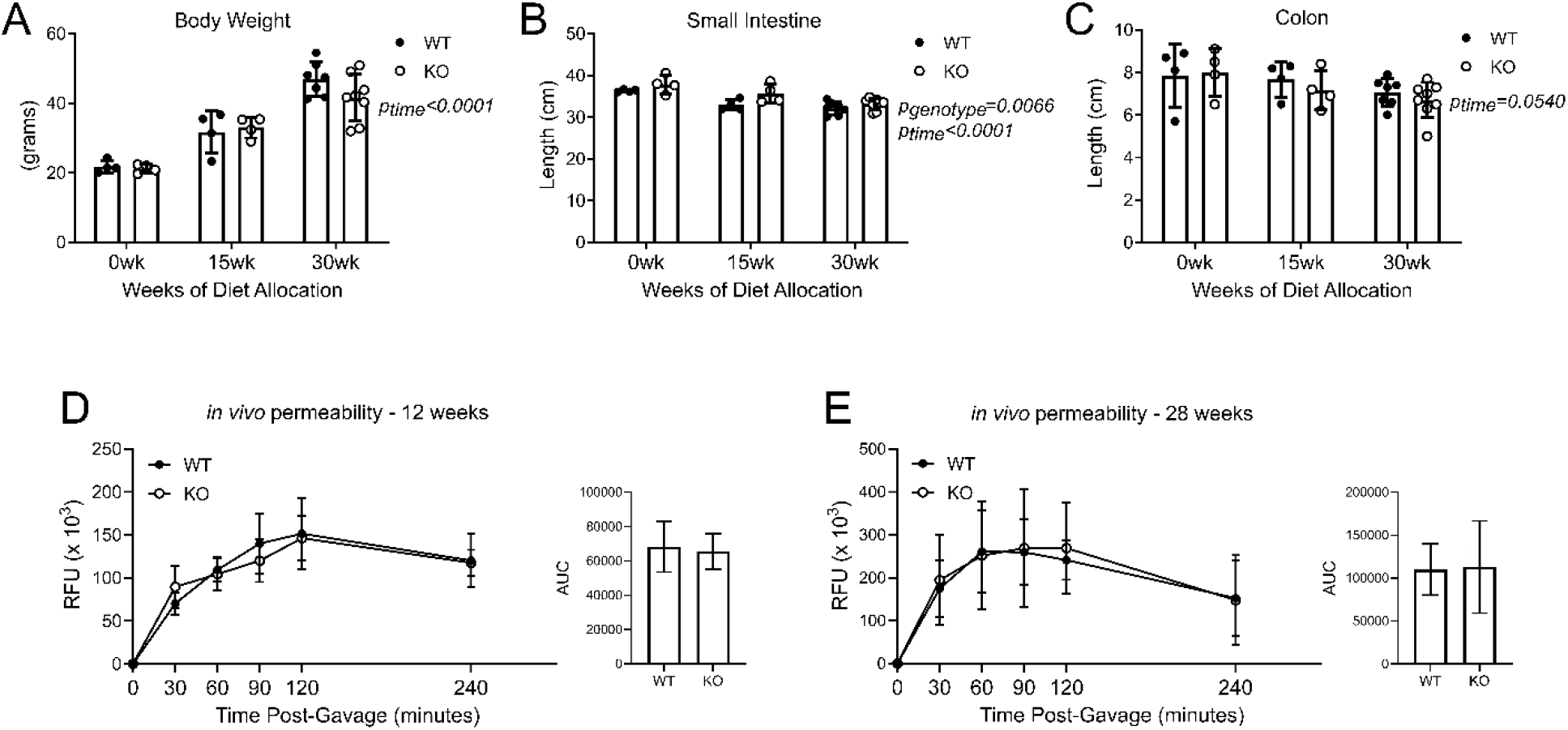
TNF does not impact body weight, intestine lengths or paracellular permeability in HF-fed female mice. Littermate wildtype (WT) and TNF^−/-^ (KO) mice were assessed at 8 weeks of age prior to diet allocation (wk0) and after allocation to a high fat diet for up to 15 weeks (wk15) or 30 weeks (wk30). (A) body weight. (B) small intestine length. (C) colon length. *In vivo* intestinal permeability to FITC-dextran after: (D) 12 weeks diet allocation, (E) 28 weeks diet allocation. Data in A-C are presented with box height at the mean and error bars at ± standard deviation. Data in D and E are presented as relative fluorescence units (RFU) with a dot at the mean ± standard deviation, with area under the curve (AUC) presented with box height at the mean with error bars indicating ± standard deviation. Data are from one independent experiment of 4-8 mice per genotype at each time point. Statistical significance was assessed by two-way ANOVA with Tukey’s post-hoc test for A-C and by two-tailed Student’s t test at each time point and AUC for D-E.

We next assessed quantities and relative prevalence of Ly6C^high^ monocytes, transitional Ly6C^+^MHCII^+^ cells, total MHCII^+^ macrophages, and monocyte-derived (CD4^−^TIM4^−^, CD4^+^TIM4^−^) and tissue-resident (CD4^+^TIM4^+^) macrophage populations in HF-fed WT and TNF^−/-^ mice by flow cytometry (Figure 6 and Supplementary Figure 6). There were no significant differences by genotype in the numbers of Ly6C^high^ monocytes (Figure 6A and 6H), transitional Ly6C^+^MHCII^+^ cells (Figure 6B and 6I), or MHCII^+^ macrophages (Figure 6C and 6J) in either the ileum or colon. The prevalence (as a proportion of total leukocytes) of Ly6C^high^ monocytes and MHCII^+^ macrophages were also similar in WT and TNF^−/-^ mice, though there was a transient increase in the proportion of Ly6C^high^ monocytes after 15 weeks in TNF^−/-^ mice (Supplementary Figure 6A-D). Quantities of monocyte-derived CD4^−^TIM4^−^ and CD4^+^TIM4^−^ macrophages, and tissue-resident CD4^+^TIM4^+^ macrophages, were not significantly different between WT and TNF^−/-^ mice at baseline, or after 15- and 30-weeks HF diet intake in either the ileum (Figure 6D-F) or the colon (Figure 6K-M). We also considered intra-macrophage population prevalence in the ileum (Figure 6G; Supplementary Figure 6E) and colon (Figure 6N; Supplementary Figure 6G). Since there were significant differences between WT and TNF^−/-^ mouse CD4^+^TIM4^−^ and CD4^+^TIM4^+^ macrophage dynamics prior to diet allocation in both tissues, we also normalized these data to the mean of the respective genotype group at baseline (Supplementary Figure 6F and 6H). While there were some changes in normalized ileum macrophage prevalence between genotypes after 30 weeks of diet intake, as described above, there were no significant differences in the quantities of cells. These data suggest that similar directional changes in WT and TNF^−/-^ intestinal monocyte-derived and tissue-resident macrophage populations occur in response to HF diet-induced obesity.

**Figure 6.**
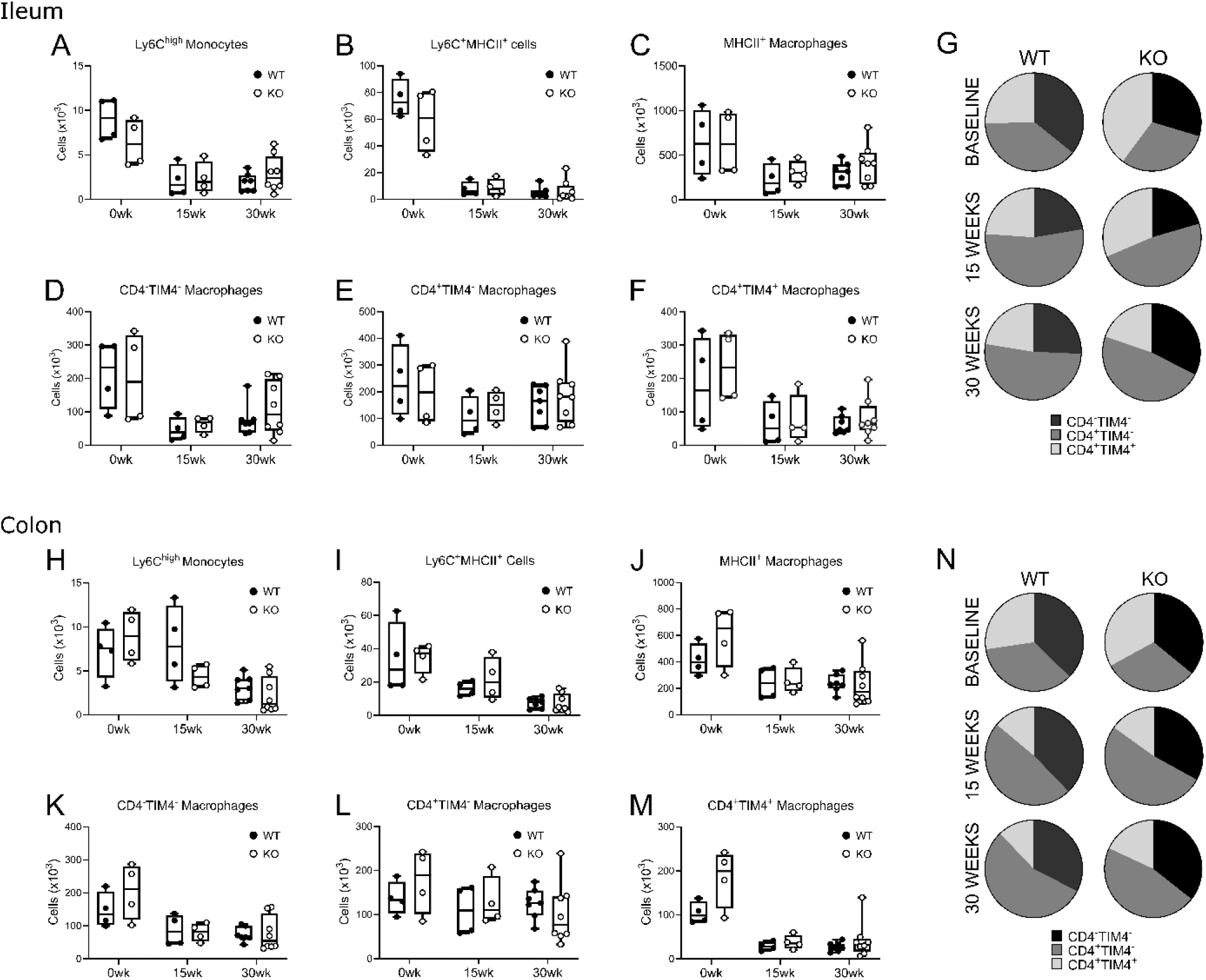
TNF does not impact effects of diet-induced obesity on intestinal monocytes and macrophages in female mice. Littermate wildtype (WT) and TNF^−/-^ (KO) mice were assessed at 8 weeks of age prior to diet allocation (wk0) and after allocation to a high fat diet. Intestinal monocyte and macrophage populations were assessed by flow cytometry in the colon and ileum. Ileum absolute cell counts (numbers) of: (A) Ly6C^high^ monocytes, (B) Ly6C^+^MHCII^+^ cells, (C) total MHCII^+^ macrophages, (D) CD4^−^TIM4^−^ macrophages, (E) CD4^+^TIM4^−^ macrophages, (F) CD4^+^TIM4^+^ macrophages. (G) summary of ileum average macrophage prevalence (as a proportion of total macrophages; also see Supplementary Figure 6). Colon absolute cell counts (numbers) of: (H) Ly6C^high^ monocytes, (I) Ly6C^+^MHCII^+^ cells, (J) total MHCII^+^ macrophages, (K) CD4^−^TIM4^−^ macrophages, (L) CD4^+^TIM4^−^ macrophages, (M) CD4^+^TIM4^+^ macrophages. (N) summary of colon average macrophage prevalence (as a proportion of total macrophages; also see Supplementary Figure 6). Each data point indicates an individual mouse. Data in A-F and H-M are presented as box and whisker plots, minimum to maximum, with the center line at the median. Data are from one independent experiment of 4-8 mice per genotype at each time point. Statistical significance was assessed by two-tailed parametric Student’s t test or Welch’s t test for unequal variances or by non-parametric Mann-Whitney U test between genotypes at each time point.

We also examined the surface phenotypes of intestinal macrophages in HF-fed WT and TNF^−/-^ mice after 15 and 30 weeks of diet intake (Figure 7). Expression of Ly6C, CD11b, CD64, CCR2, F4/80, and MHCII on ileum macrophages was similar between TNF^−/-^ and WT mice after 15 or 30 weeks of HF diet feeding. Colon CD4^+^TIM4^+^ macrophages in HF-fed TNF^−/-^ mice compared to WT mice had decreased expression of CD11b after 15 weeks (Supplementary Figure 7J), and decreased expression of CD64 after 30 weeks (Figure 7K), but there were no other significant differences between genotypes. Together our data suggest that in female mice, the absence of TNF does not prevent changes to body weight, intestinal length, permeability, nor intestinal macrophage numbers or surface phenotype that accompany HF diet-induced obesity.

**Figure 7.**
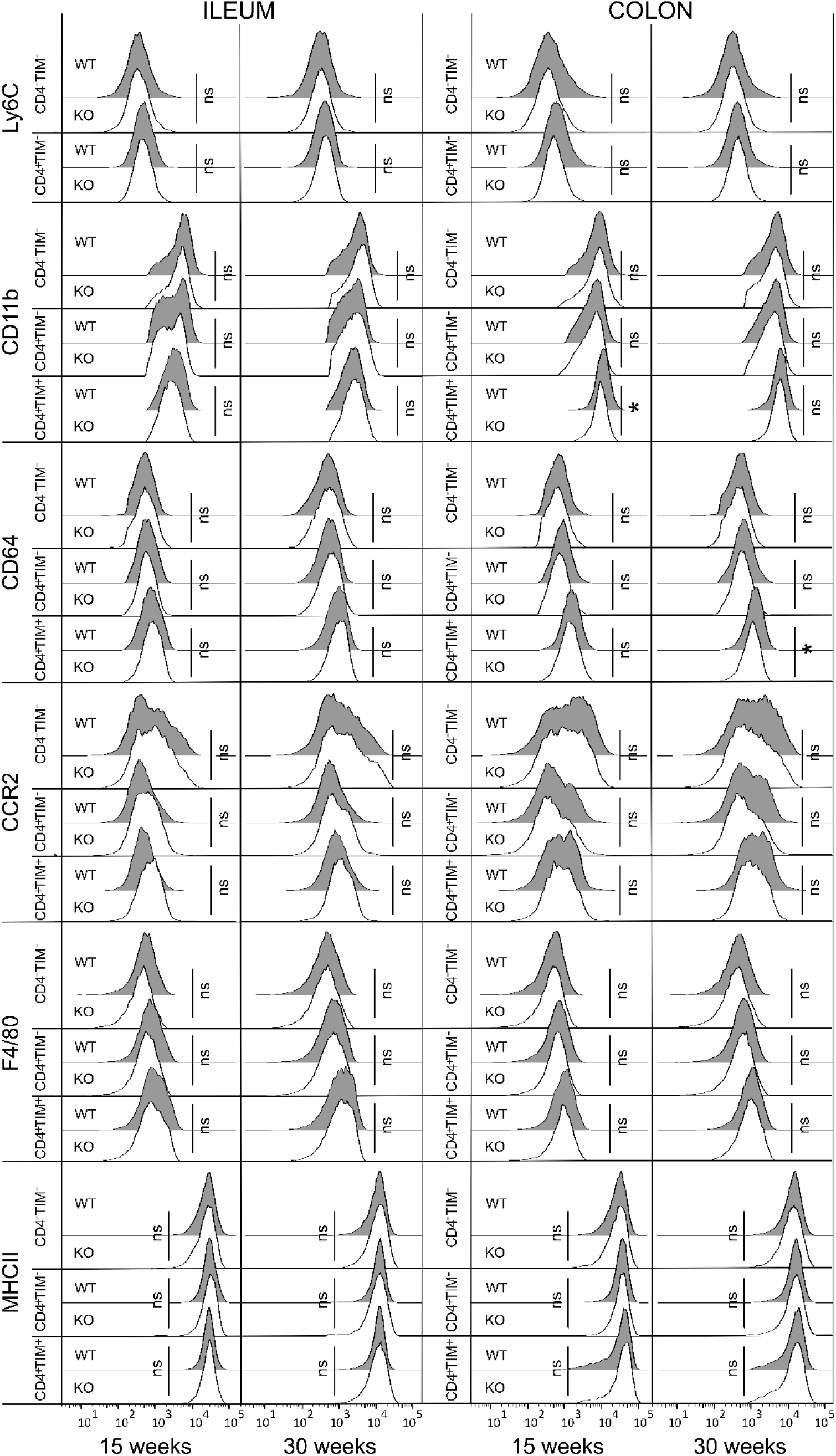
TNF does not mediate changes to macrophage surface phenotype in HF-fed female mice. Flow cytometry analysis of CD4^−^TIM4^−^, CD4^+^TIM4^−^, and CD4^+^TIM4^+^ macrophage populations in the ileums and colons of littermate wildtype (WT) and TNF^−/-^ (KO) female mice on high fat diet for 15 or 30 weeks. Macrophage phenotype was assessed by examining surface expression of Ly6C, CD11b, CD64, CCR2, F4/80, and MHCII. Data are reported as geometric mean fluorescence intensity and are visualized by concatenating uncompensated events in FlowJo for each mouse and indicated macrophage population at each time point grouped by genotype, and then geometric mean fluorescence intensity expression data of each concatenated group was overlaid onto the same histogram plot. Data are from one experiment of WT n=4 and KO n=4 at 15 weeks, and WT n=7 and KO n=8 at 30 weeks. Statistical significance was assessed by two- tailed parametric Student’s t test or Welch’s t test for unequal variances or by non-parametric Mann-Whitney U test between macrophage populations by genotype at each time point. *\*p<0*.*05*.

## Discussion

In this study we found that diet-induced obesity in female mice has early and sustained effects on intestinal monocytes and macrophages, including a decrease in monocyte recruitment, and depletion of monocyte-derived macrophages as well as tissue-resident macrophages, without a subsequent compensatory increase in local macrophage proliferation. Our data confirm that the recruitment, phenotype, and functions of intestinal monocytes and macrophages are affected by changes within their local tissue microenvironment (Grainger *et al*., 2017), and support previous observations of altered intestinal macrophage populations in obese humans (Monteiro-Sepulveda *et al*., 2015; Rohm *et al*., 2021). Our data contrast with reports of accumulation of pro-inflammatory TNF-producing monocyte-derived macrophages in metabolic tissues in diet-induced obesity (Weisberg *et al*., 2003; Kanda *et al*., 2006), as well as within the intestines in response to acute or chronic inflammation from infection or colitis (Zigmond *et al*., 2012; Bain *et al*., 2013; Bain *et al*., 2018). Nonetheless, our data are consistent with histological reports of low-grade inflammation in the intestines compared with metabolic tissues in obesity (Johnson *et al*., 2015; Luck *et al*., 2015), as well as with observations of the ability of intestinal macrophages to remain hypo-responsive to exogenous stimulation (Grainger *et al*., 2017; Muller *et al*., 2020). Therefore, our findings are reflective of the intrinsically anti-inflammatory and tolerogenic environment of the gut (Greenwood-Van Meerveld *et al*., 2017). While outside the scope of this study, further research that incorporates *in situ* immunohistochemistry or immunofluorescence techniques would provide further insight into any spatially-restricted changes in intestinal macrophage populations. It is also possible that there are pro-inflammatory shifts in other intestinal immune cells in response to diet-induced obesity, as has been reported for some T cell populations (Garidou *et al*., 2015; Luck *et al*., 2015; Monteiro-Sepulveda *et al*., 2015), which may influence macrophage composition and function.

We found that HF diet-induced obesity results in tissue- and time-dependent physiological changes within the intestines, including shortening of ileum and colon lengths, increased colon ion transport, and elevated paracellular permeability. These data support prior literature showing that diet-induced obesity impairs maintenance of the intestinal epithelium (Cani *et al*., 2007; Cani *et al*., 2008; Mah *et al*., 2014; Johnson *et al*., 2015; Kawano *et al*., 2016). Diet-induced obesity has also been reported to dysregulate blood vessel barriers (Mouries *et al*., 2019), enteric neuron activity (Stenkamp-Strahm *et al*., 2013), and gastrointestinal motility (Mushref & Srinivasan, 2013). As all of these functions are affected by disruption of monocyte recruitment and/or macrophage depletion (Muller *et al*., 2014; Gabanyi *et al*., 2016; De Schepper *et al*., 2018; Honda *et al*., 2020), an obesity-associated reduction in intestinal monocyte and macrophage numbers, as observed in this study, may contribute to those physiological changes. Furthermore, as mentioned, under homeostatic conditions TNF regulates intestinal epithelial barrier function (Delgado & Brunner, 2019). While overproduction of TNF is typically associated with damage to the intestinal epithelium, it has been demonstrated that tissue-resident macrophage production of TNF is actually protective against colitis induced by disruption of the intestinal epithelium (Kaya *et al*., 2020). Our data suggest that intestinal macrophages do not contribute to local inflammation via TNF in diet-induced obesity, yet the decrease in relative TNF production by both ileum and colon macrophages that we found in HF-fed mice may have contributed to the observed increases in paracellular permeability. Thus, obesity-associated loss of intestinal homeostasis is likely mediated by physiological changes that contribute to alterations in local macrophage populations, and *vice versa*.

Obesity-associated changes to the microbiota and microbial metabolite production may also influence local macrophage populations (Caprara *et al*., 2020). In addition, it should be noted that saturated fatty acids found in high fat diets can induce macrophage mitochondrial dysfunction and apoptosis (Martins de Lima *et al*., 2006; Li *et al*., 2018). Exacerbated pathology has been reported after high fat diet feeding in mouse models of colitis in the absence of obesity (Kim *et al*., 2010; Gruber *et al*., 2013). Most dietary fats are absorbed in the small intestine (Greenwood-Van Meerveld *et al*., 2017), which may explain why we observed a greater and more sustained depletion of monocyte-derived macrophages and increase in paracellular permeability in the ileum compared to the colon. Given the ubiquitous use of high fat diets in mouse models of obesity, consideration of independent and combined effects of diet and obesity on intestinal macrophages and other immune cells merits further research. Irrespective, our data align with previous work showing that diet-induced obesity results in loss of intestinal homeostasis and barrier function (Cani *et al*., 2008; Kawano *et al*., 2016; Luck *et al*., 2019), which may promote translocation of luminal contents including bacterial products into circulation that contribute to the gradual development of systemic inflammation and metabolic dysfunction.

The gut is a dynamic organ, which experiences natural changes in structure, function, and immune cell composition throughout the life course (Walrath *et al*., 2021). In young chow-fed mice, we observed a gradual increase in the prevalence of monocyte-derived macrophages and an accompanying decrease in the prevalence of tissue-resident macrophages within ileum and colon tissues between the short, intermediate, and prolonged time points of assessment between 6 and 18 weeks. These data are consistent with previous reports of macrophage population composition and turnover, showing that there is a gradual increase in monocyte-derived macrophages in intestinal (and other) tissues from birth in mice (Shaw *et al*., 2018; Liu *et al*., 2019) and in humans (Bernardo *et al*., 2018; Bujko *et al*., 2018; Wright *et al*., 2021).

Accordingly, the disruptions in monocyte-derived and tissue-resident macrophage turnover dynamics that we observed due to HF diet-induced obesity could have more severe long-term effects on intestinal function if they occur earlier in life than in mature adults (i.e., prior to stabilization of intestinal macrophage populations). Further studies should consider if effects of diet-induced obesity on intestinal macrophages differ in contexts that impact intestinal function and permeability, including pregnancy and lactation (Hammond, 1997), increasing age (DeJong *et al*., 2020), and when homeostasis is already disturbed, such as in response to acute or chronic inflammation (Eckmann & Kagnoff, 2005; Yan *et al*., 2009).

Our finding that TNF did not contribute to changes in intestinal physiology and macrophages in HF diet-induced obesity was surprising, because previous research suggests that TNF contributes to intestinal inflammation in diet-induced obesity (Ding *et al*., 2010; Lam *et al*., 2012; Liu *et al*., 2012; Kawano *et al*., 2016). Yet, there are conflicting data from animal models and clinical reports on TNF and anti-TNF therapies, implying that TNF has pleiotropic effects in acute and chronic inflammation within the intestines. For example, TNF^−/-^ mice are not susceptible to acute or chronic TNBS-induced colitis (Noti *et al*., 2010), but are susceptible to acute DSS-induced colitis (Kojouharoff *et al*., 1997). As well, anti-TNF therapy leads to worse tissue damage in acute DSS-colitis (Naito *et al*., 2003), but reduces inflammation in chronic colitis (Kojouharoff *et al*., 1997). Treatment failure of anti-TNF therapy is documented in inflammatory bowel disease, with both primary non-response or loss of response after initial successful treatment (Fine *et al*., 2019), which suggests that there may be different etiologies of intestinal inflammation that are not TNF-dependent. Collectively, these data caution that context may dictate whether TNF has beneficial or deleterious effects. These effects could in addition be further modified by biological sex, which influences both immunity and metabolism (Tramunt *et al*., 2020; Breznik *et al*., 2021b). It has been reported that TNF-producing monocyte-derived colon macrophages contribute to obesity-associated metabolic dysfunction in male mice (Kawano *et al*., 2016; Rohm *et al*., 2022). However, we have observed that while both obese male and female mice have an increase in circulating inflammatory TNF-producing Ly6C^high^ monocytes and adipose tissue macrophage accumulation, in female mice the metabolic dysregulation is attenuated, and these changes are not dependent on TNF (Breznik *et al*., 2018; Breznik *et al*., 2021b). While turnover of intestinal macrophages has been reported to be similar in non-obese young male and female mice (Bain *et al*., 2016; Liu *et al*., 2019), it has been found that estrogen treatment has anti-inflammatory effects in mouse models of colitis (Verdú *et al*., 2002; Bábíčková *et al*., 2015), and moreover, estrogen may attenuate effects of diet-induced obesity in the colon (Hases *et al*., 2020). *A priori* consideration of biological sex and hormones may therefore be indispensable in disentangling the complex effects of TNF in both acute and chronic intestinal inflammation. Future investigations should be designed to evaluate how estrogens and androgens modify the role of TNF in mediating changes to intestinal macrophage populations and maintenance of the intestinal barrier in obesity.

In conclusion, we have identified tissue-specific effects of diet-induced obesity on monocyte-derived and tissue-resident intestinal macrophage numbers, prevalence, proliferation, surface phenotype, and cytokine profiles. These data emphasize the importance of studying macrophages within their local tissue environment, and contribute to the growing evidence that biological sex influences cellular immune responses to obesity-associated chronic inflammation.

## Data availability

Data from this study are available from the corresponding author upon reasonable request.

## Acknowledgements

The authors would like to thank John Grainger for his guidance on tissue processing and flow cytometry protocol design, Tatiane Ribeiro for her guidance on *in vivo* permeability assays, as well as Erica Yeo, Erica DeJong, Janine Strehmel, Melodie Kim, Brianna Kennelly, and Braeden Cowbrough for technical support, Hong Liang and the McMaster Immunology Research Centre flow cytometry facility, and Erin Demask and Tina Walker for provision of media. The graphical abstract was created using BioRender.com.

