## SupplementaryFiles for "Diet-induced obesity alters intestinal monocyte-derived and tissue-resident macrophages in female mice independent of TNF"

**Supplementary Table 1. Antibodies for intestinal immune cell surface stain**

| Antibody | Fluorophore | Company and Catalogue # | Dilution |
|---|---|---|---|
| Ly6C | Alexa Fluor 488 | BioLegend 128022 (HK1.4) | 1:100 |
| Tim-4 | PE | BioLegend 130005 (RMT4-54) | 1:50 |
| CD64 | PE-Dazzle 594 | BioLegend 13919 (X54-5/7.1) | 1:200 |
| CD11b | PerCP-Cy5.5 | eBioscience 45-0112-82 (M1/70) | 1:200 |
| CCR2 | Brilliant Violet 421 | BioLegend 150605 (SA203G11) | 1:100 |
| CD45 | Brilliant Violet 510 | BioLegend 103137 (30-F11) | 1:200 |
| F4/80 | Brilliant Violet 605 | BioLegend 123133 (BM8) | 1:100 |
| CD4 | APC | BioLegend 100515 (RM4-5) | 1:50 |
| MHC II | AF700 | eBioscience 56-5231-80 (M5/114.15.2) | 1:200 |
| Live-Dead NIR | APC-Cy7 | Invitrogen L34975 | 1:200 |
| CD3 | PE-Cy7 | eBioscience 25-0031-81 (145-2C11) | 1:200 |
| B220 | PE-Cy7 | eBioscience 25-0452-81 (RA3-6B2) | 1:200 |
| Ly6G | PE-Cy7 | eBioscience 25-0112-81 (M1/70) | 1:200 |

**Supplementary Table 2. Antibodies for intestinal immune cell surface stain (to be followed by intracellular staining)**

| Antibody | Fluorophore | Company and Catalogue # | Dilution |
|---|---|---|---|
| Tim-4 | PE | BioLegend 130005 (RMT4-54) | 1:50 |
| CD64 | PE-Dazzle 594 | BioLegend 13919 (X54-5/7.1) | 1:200 |
| CD11b | PE-Cy7 | eBioscience 25-0112-81 (M1/70) | 1:200 |
| CD45 | Brilliant Violet 510 | BioLegend 103137 (30-F11) | 1:200 |
| CD4 | APC | BioLegend 100515 (RM4-5) | 1:50 |
| MHC II | AF700 | eBioscience 56-5231-80 (M5/114.15.2) | 1:200 |
| Live-Dead NIR | APC-Cy7 | Invitrogen L34975 | 1:200 |

**Supplementary Table 3. Antibodies for intracellular intestinal immune cell staining**

| Antibody | Fluorophore | Company and Catalogue # | Dilution |
|---|---|---|---|
| TNF | Alexa Fluor 488 | eBioscience 53-7321-82 (MP6-XT22) | 1:67 |
| IL-10 | PerCP-Cy5.5 | eBioscience 45-7101-80 (JES5-16E3) | 1:67 |
| Ki67 | Brilliant Violet 605 | BioLegend 652413 (16A8) | 1:67 |

**Supplementary Figure 1. Flow cytometry gating to identify intestinal macrophages.**

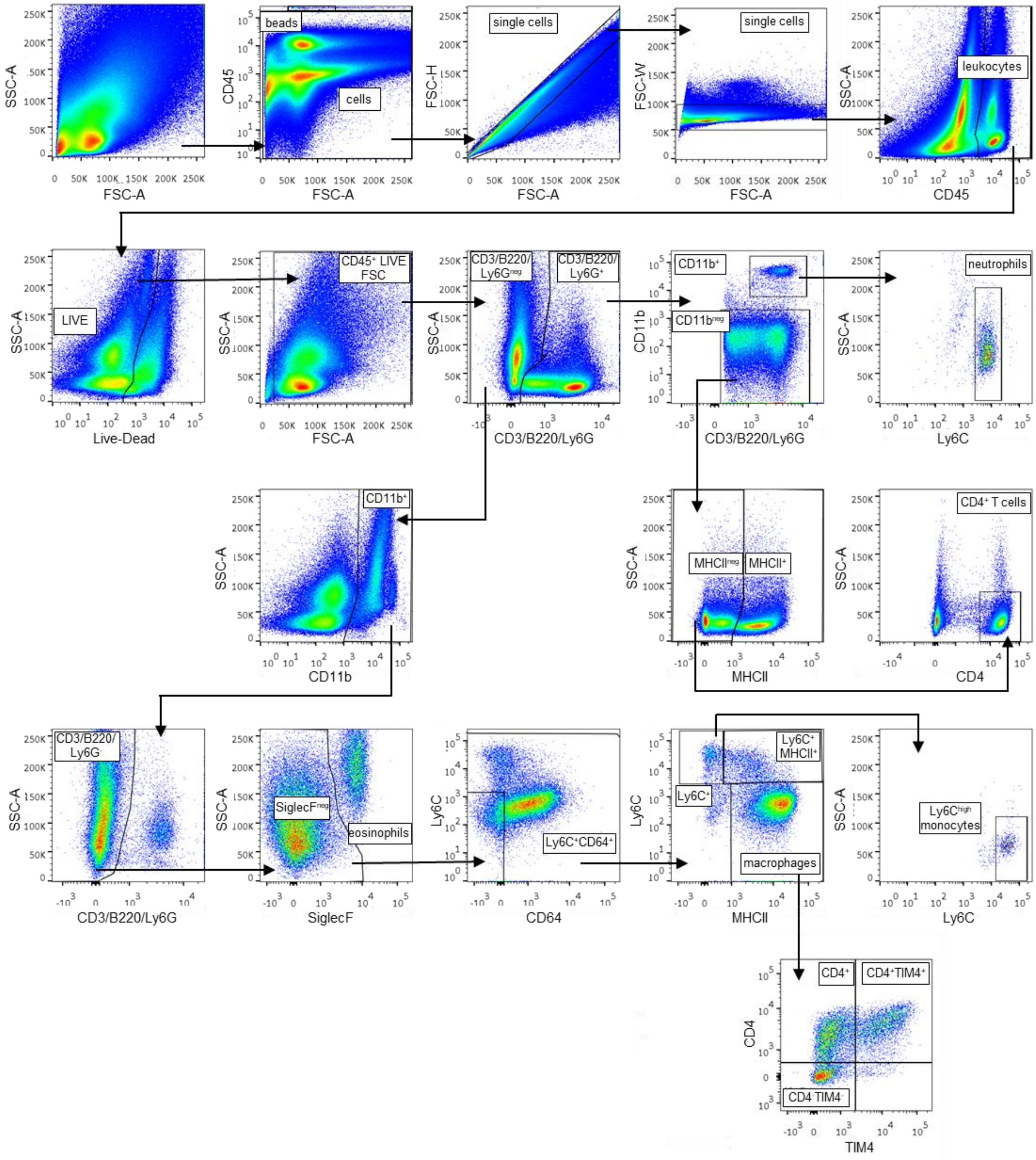

**Supplementary Figure 2. Flow cytometry gating for macrophage intracellular staining.**

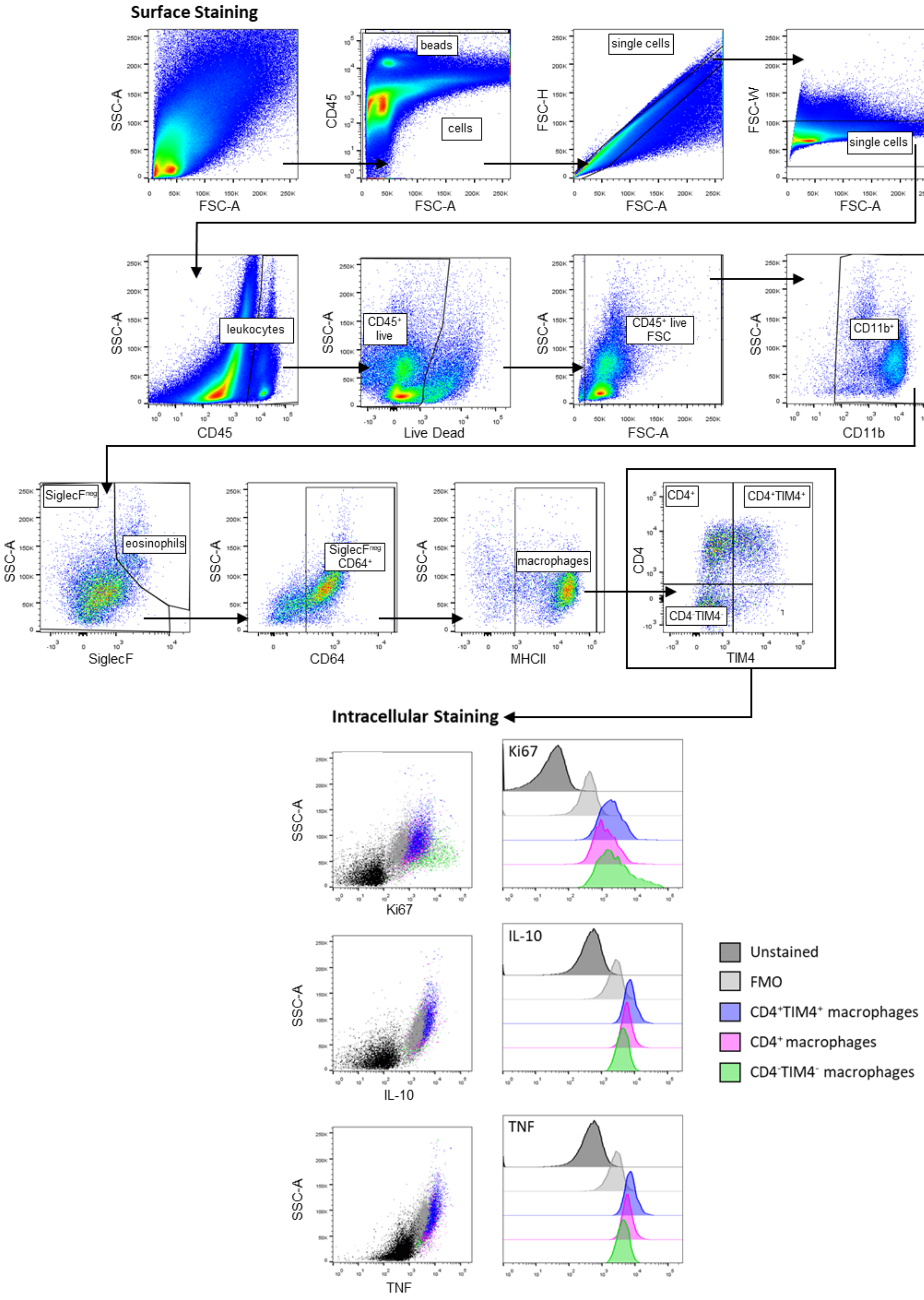

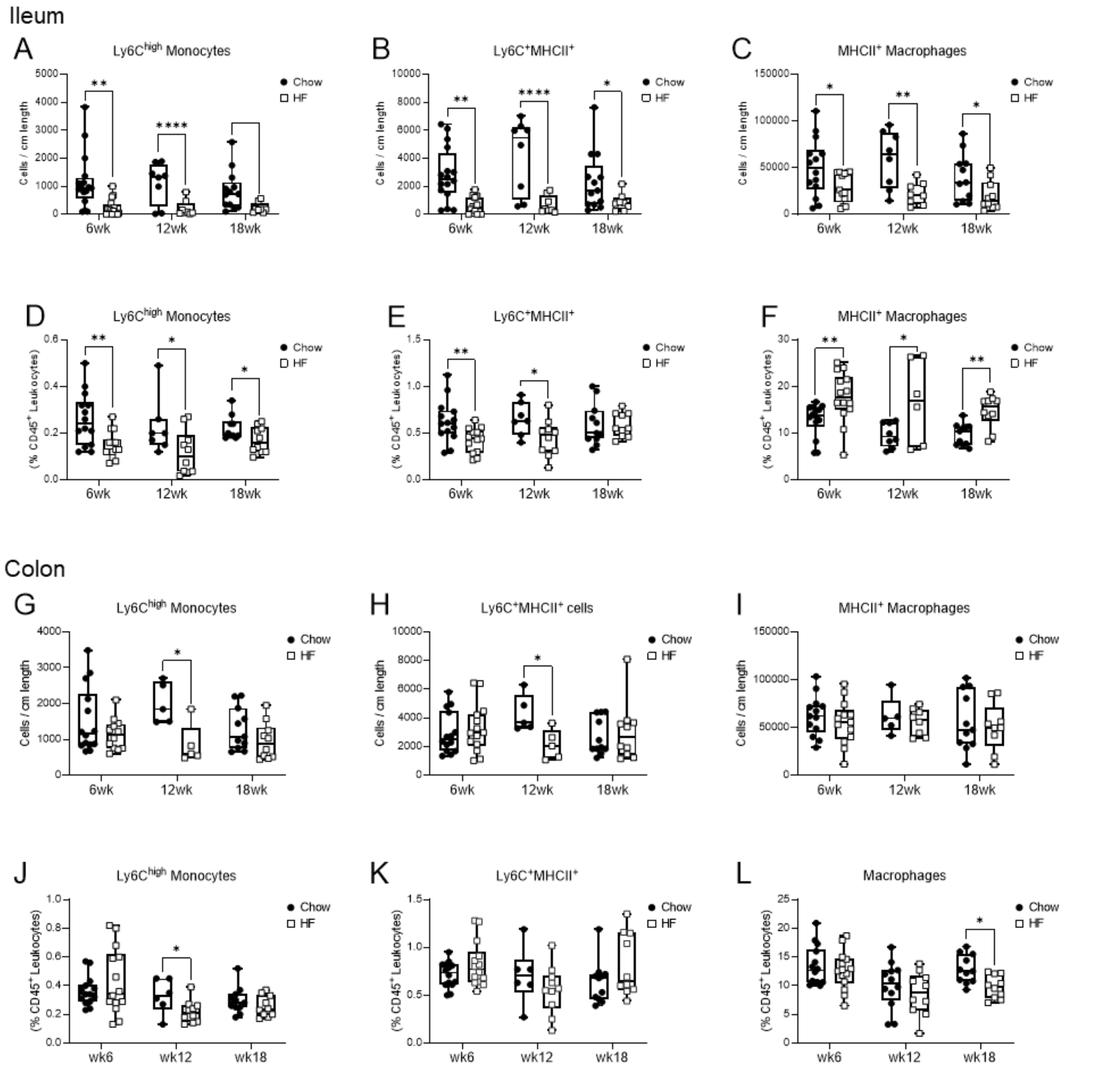


**Supplementary Figure 3. Intestinal monocyte and macrophage cell numbers per tissue length and prevalence in chow and HF-fed mice.**

Intestinal monocyte and macrophage populations were assessed by flow cytometry in the colon and ileum after 6, 12, or 18 weeks of diet allocation to standard chow (Chow) or high fat (HF) diet. Cell numbers adjusted by ileum tissue length of: (A) $CD4^{-}TIM4^{-}$ macrophages, (B) $CD4^{+}$ macrophages, (C) $CD4^{+}TIM4^{+}$ macrophages. Ileum prevalence (as a proportion of total $CD45^{+}$ leukocytes) of: (D) $Ly6C^{high}$ monocytes, (E) $Ly6C^{+}MHCII^{+}$ cells, (F) total $MHCII^{+}$ macrophages. Cell numbers adjusted by colon tissue length of: (G) $CD4^{-}TIM4^{-}$ macrophages, (H) $CD4^{+}$ macrophages, (I) $CD4^{+}TIM4^{+}$ macrophages. Colon prevalence (as a proportion of total $CD45^{+}$ leukocytes) of: (J) $Ly6C^{high}$ monocytes, (K) $Ly6C^{+}MHCII^{+}$ cells, (L) total $MHCII^{+}$ macrophages. Each data point indicates an individual mouse. Data are presented as box and whisker plots, minimum to maximum, where the center line indicates the median. Data are pooled from two to three independent experiments of n=4-5 mice per group. Statistical significance was assessed by two-tailed parametric Student's t test or Welch's t test for unequal variances or by non-parametric Mann-Whitney U test between macrophage populations by diet at each time point. *$p<0.05$, **$p<0.01$, ****$p<0.0001$.*

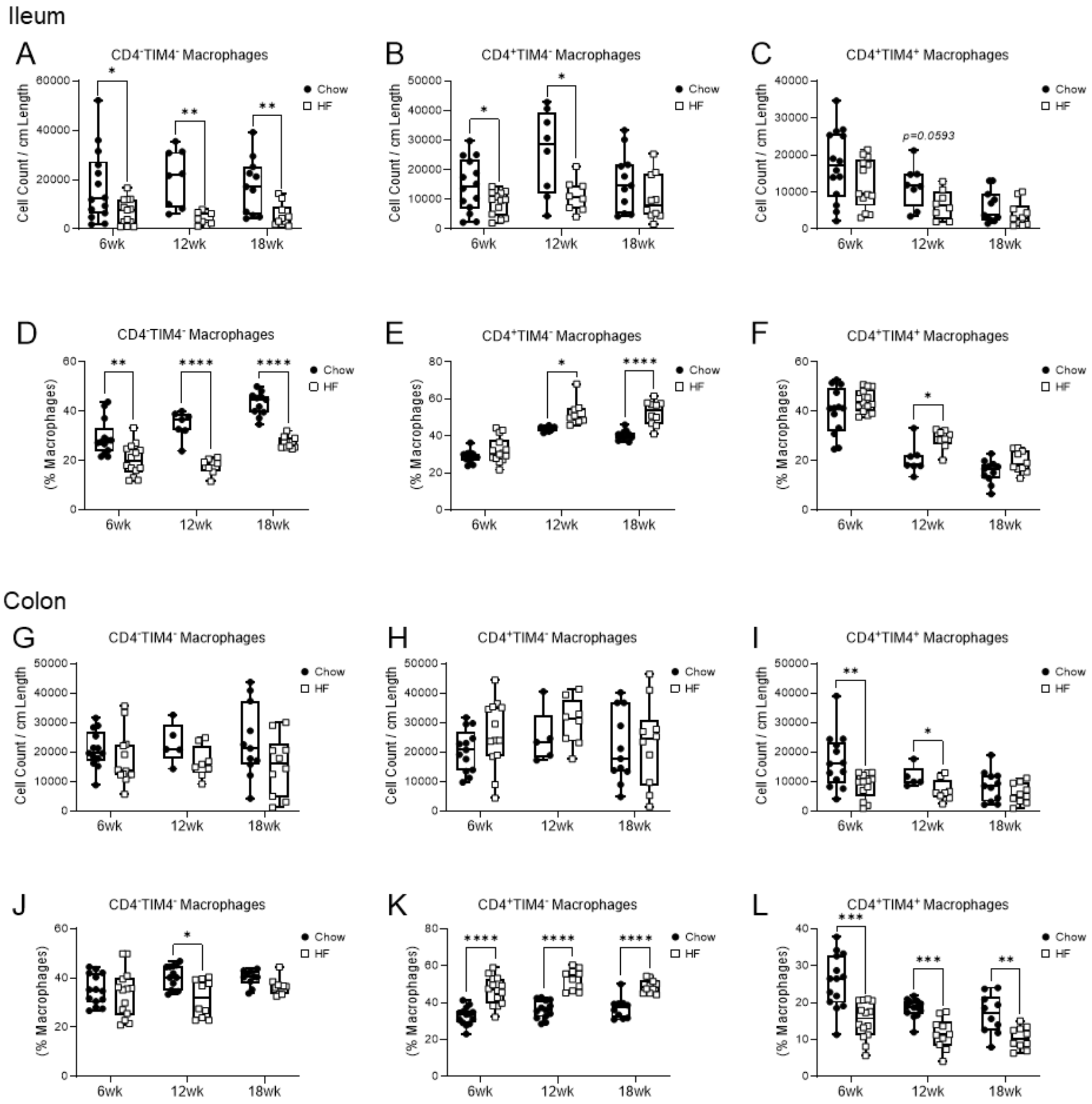


**Supplementary Figure 4. Intestinal macrophage cell numbers per tissue length and prevalence in chow and HF-fed mice.**

Intestinal monocyte and macrophage populations were assessed by flow cytometry in the colon and ileum after 6, 12, or 18 weeks of diet allocation to standard chow (Chow) or 60% high fat (HF) diet. Cell numbers adjusted by ileum tissue length of: (A) $CD4^{-}TIM4^{-}$ macrophages, (B) $CD4^{+}$ macrophages, (C) $CD4^{+}TIM4^{+}$ macrophages. $CD4^{-}TIM4^{-}$, $CD4^{+}$, and $CD4^{+}TIM4^{+}$ macrophage population prevalence in the ileum after: (D) 6 weeks, (E) 12 weeks, (F) 18 weeks diet allocation. Cell numbers adjusted by colon tissue length of: (G) $CD4^{-}TIM4^{-}$ macrophages, (H) $CD4^{+}$ macrophages, (I) $CD4^{+}TIM4^{+}$ macrophages. $CD4^{-}TIM4^{-}$, $CD4^{+}$, and $CD4^{+}TIM4^{+}$ macrophage population prevalence in the colon after: (J) 6 weeks, (K) 12 weeks, (L) 18 weeks diet allocation. Each data point indicates an individual mouse. Data are presented as box and whisker plots, minimum to maximum, where the center line indicates the median. Data are pooled from two to three independent experiments of n=4-5 mice per group. Statistical significance was assessed by two-tailed parametric Student's t test or Welch's t test for unequal variances or by non-parametric Mann-Whitney U test between macrophage populations by diet at each time point. **p<0.05, **p<0.01, ***p<0.001, ****p<0.0001.*

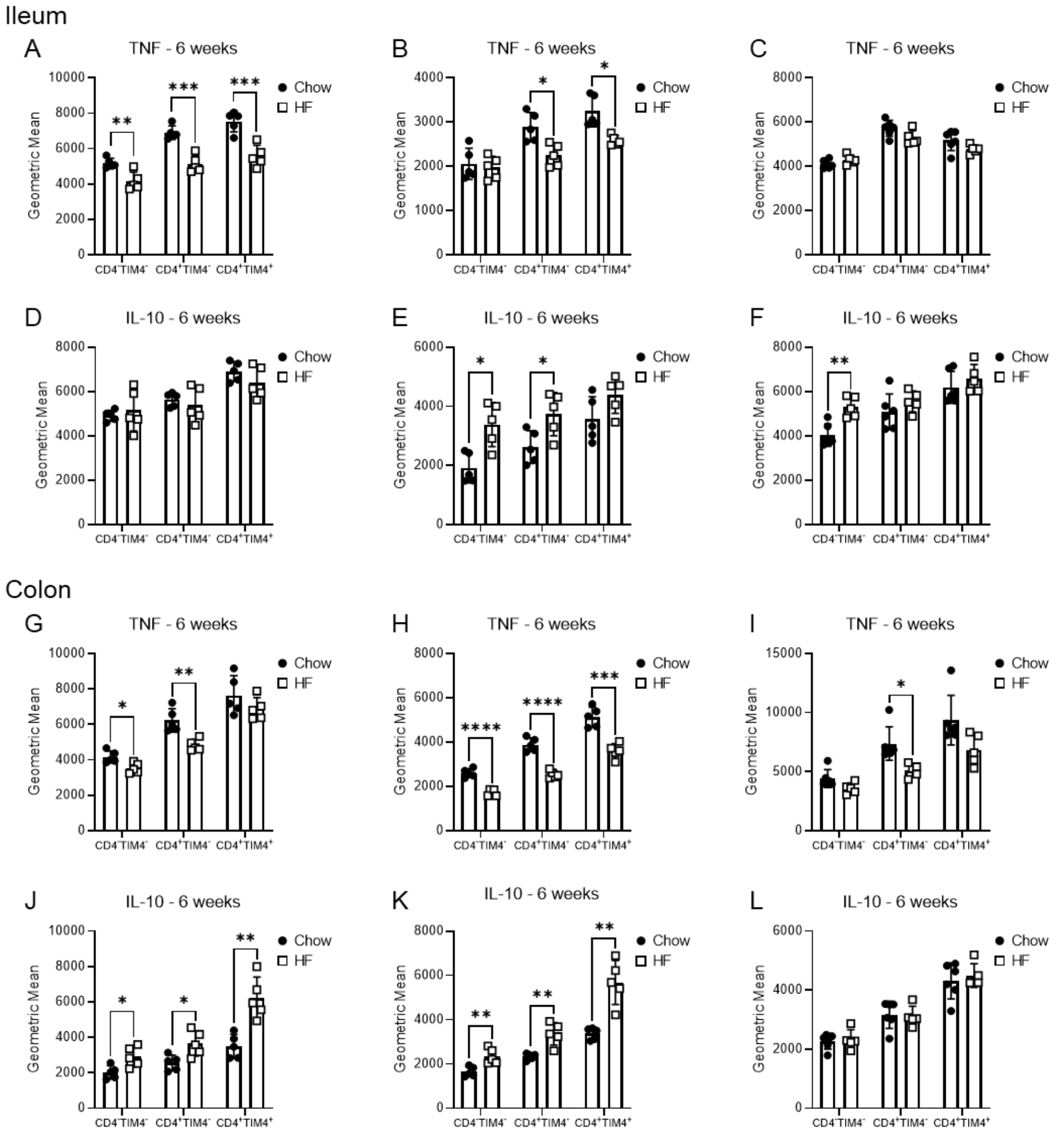


**Supplementary Figure 5. TNF and IL-10 expression in ileum and colon macrophages of chow and HF-fed mice.**

Ileum and colon $CD4^-TIM4^-$, $CD4^+$, and $CD4^+TIM4^+$ macrophage intracellular expression of TNF and IL-10 was assessed by flow cytometry after 6, 12, or 18 weeks of allocation to standard control chow (Chow) or 60% high fat (HF) diet. Ileum macrophage intracellular expression of TNF after: (A) 6 weeks, (B) 12 weeks, (C) 18 weeks. Ileum macrophage intracellular expression of IL-10 after: (D) 6 weeks, (E) 12 weeks, (F) 18 weeks. Colon macrophage intracellular expression of TNF after: (G) 6 weeks, (H) 12 weeks, (I) 18 weeks. Colon macrophage intracellular expression of IL-10 after: (J) 6 weeks, (K) 12 weeks, (L) 18 weeks. Each data point indicates an individual mouse. Data are from one independent experiment at each time point of n=5-6 mice per group. Data are presented as box and whisker plots, minimum to maximum, where the center line indicates the median. Statistical significance was assessed by two-tailed parametric Student's t test or Welch's t test for unequal variances or by non-parametric Mann-Whitney U test between macrophage populations by diet at each time point. *$*p<0.05$, $**p<0.01$, $***p<0.001$, $****p<0.0001$.*

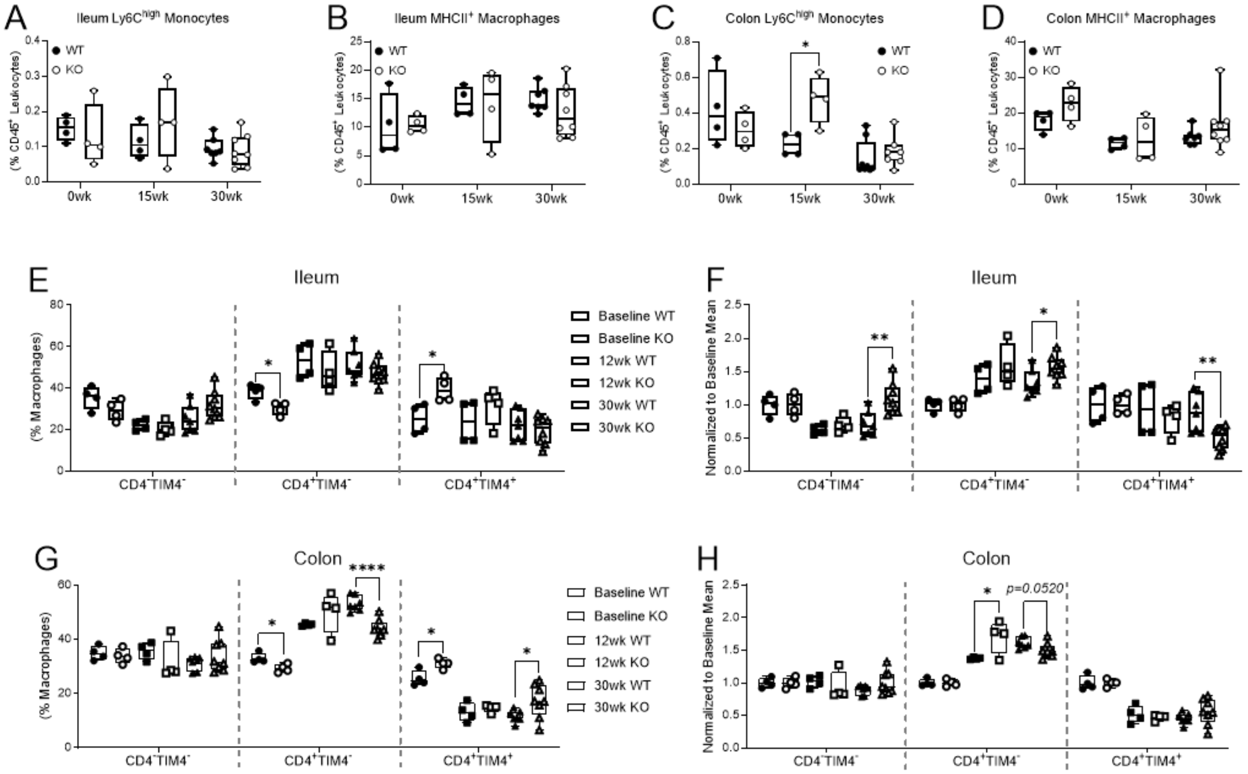


**Supplementary Figure 6. Intestinal monocyte and macrophage prevalence in HF-fed littermate WT and TNF$^{-/-}$ mice.**

Littermate wildtype (WT) and TNF$^{-/-}$ (KO) mouse intestinal monocytes and macrophages were assessed by flow cytometry at 8 weeks of age (0wk/Baseline), and after 15 weeks (15wk) or 30 weeks (30wk) of high fat diet allocation. Ileum prevalence (as a proportion of total CD45$^{+}$ leukocytes) of: (A) Ly6C$^{high}$ monocytes, (B) total MHCII$^{+}$ macrophages. Colon prevalence (as a proportion of total CD45$^{+}$ leukocytes) of: (C) Ly6C$^{high}$ monocytes, (D) total MHCII$^{+}$ macrophages. (E) ileum prevalence of CD4$^{-}$TIM4$^{-}$, CD4$^{+}$, and CD4$^{+}$TIM4$^{+}$ macrophages (as a proportion of total macrophages). (F) ileum prevalence of CD4$^{-}$TIM4$^{-}$, CD4$^{+}$, and CD4$^{+}$TIM4$^{+}$ macrophages normalized to the mean of the baseline data to adjust for between-genotype differences. (G) colon prevalence of CD4$^{-}$TIM4$^{-}$, CD4$^{+}$, and CD4$^{+}$TIM4$^{+}$ macrophages (as a proportion of total macrophages). (F) colon prevalence of CD4$^{-}$TIM4$^{-}$, CD4$^{+}$, and CD4$^{+}$TIM4$^{+}$ macrophages normalized to the mean of the baseline data to adjust for between-genotype differences by dividing each data point from 15- and 30-weeks diet intake to the mean of the respective genotype group at baseline. Each data point indicates an individual mouse. Data are presented as box and whisker plots, minimum to maximum, with the center line at the median. Data are from one independent experiment of 4-8 mice per genotype at each time point. Statistical significance was assessed by two-tailed parametric Student's t test or Welch's t test for unequal variances or by non-parametric Mann-Whitney U test between macrophage populations by genotype at each time point. *$p<0.05$, **$p<0.01$, ****$p<0.0001$.*
